# *afila*, the origin and nature of a major innovation in the history of pea breeding

**DOI:** 10.1101/2023.07.19.549624

**Authors:** Nadim Tayeh, Julie Hofer, Grégoire Aubert, Françoise Jacquin, Lynda Turner, Jonathan Kreplak, Pirita Paajanen, Christine Le Signor, Marion Dalmais, Stéphanie Pflieger, Valérie Geffroy, Noel Ellis, Judith Burstin

## Abstract

The *afila* (*af*) mutation of *Pisum sativum* L. (pea) is characterised by leaves that are composed of a basal pair of stipules, a petiole and a branched mass of tendrils. These are bipinnate leaves in which the leaflet primordia are replaced by midrib-like, or terminal tendril, primordia. The phenotype was first reported as a spontaneous mutation in 1953, and several reports of spontaneously occurring *af* mutants and induced mutations have been published since then. Despite its wide-scale introgression to improve standing ability in combine-harvested dry pea crops, the molecular basis of *af* has remained unknown. Here, we show that the deletion of two tandemly-arrayed Q-type Cys(2)His(2)-zinc finger transcription factors, viz. *PsPALM1a* and *PsPALM1b*, is responsible for the af phenotype. Based on molecular evidence for the presence/absence of seven consecutive pea genes, we identified eight haplotypes in the genomic region of chromosome 2 that harbours *af*. These haplotypes differ in the presence or absence of *PsPALM1a-b* and close genes and in the size of the deletion. Representative cultivars and spontaneous or induced mutants were assigned to the different haplotypes. The hitherto unrecognised diversity at the *af* locus reveals highly rich, unexplored, potential for pea improvement and sheds light on the breeding history of pea. This knowledge can also be used to breed innovative cultivars in related crops.

## Introduction

Plant breeding exploits natural or induced genetic variability as the source of superior alleles in progenitor lines. Selection for desired phenotypes relies on family-based or marker-based predictions. Several outstanding crop-improvement successes have been achieved by the modification of plant architecture, as exemplified by the introduction of dwarfing genes in cereals (Borlaug 1983; Lumpkin 2015). However, in most cases, the molecular determinants responsible for increased genetic gain are unknown. Even though it is not required for the breeding process *per se*, the identification of such determinants is important to minimize linkage drag around target loci (Collard and Mackill 2008) and to search for useful variant alleles. Such step enables wiser and more effective use of the genetic effects at the intra-specific level and also at the inter-specific level through translational approaches (Jacob et al. 2018).

Pea (*Pisum sativum* L.) is an important crop for food and feed production, because of its protein-rich dry seeds, and plays a strategic role in the sustainable intensification of agriculture to provide food security (McDermott and Wyatt 2017; Poore and Nemecek 2018). Since the mid-1970s, pea has been developed as a high-quality protein crop in Europe (Snoad 1981), in part to reduce over-dependence upon imported soya (de Visser et al. 2014). Pea is a cool-season legume belonging to the Fabeae tribe of the inverted repeat-lacking clade (IRLC) of the Papilionoideae subfamily of legumes (Azani et al. 2017). It was a model organism of choice to study the basic principles of inheritance and independent segregation (Knight 1799; Mendel 1866) and to study plant morphology and physiology (Dostál 1941; Murfet 1977; Smýkal 2014).

Leaves are usually flat organs, that account for the majority of the photosynthetic activity and carbon supply required during the life cycle of a plant (Schiltz et al. 2005). Leaf shape and structure affect photosynthesis and transpiration and are critical for the energy efficiency and water economy of crop plants. Leaf form also contributes to the microclimate of the crop canopy and thus influence disease progression (Schoeny et al. 2008). Genes that control the size and shape of leaves are therefore of primary importance to agriculture. Wild type peas have pinnately compound leaves; a mature adult leaf consists of a basal pair of stipules, proximal leaflet pairs, distal tendril pairs, and a terminal tendril (Figure 1A; Hofer and Ellis 1998; Gourlay et al. 2000). The molecular bases of different mutations affecting leaf morphogenesis in pea have been described, including *unifoliata* (*uni*; Eriksson 1929; Hofer et al. 1997), *stamina-pistilloida* (*stp*; Monti and Devreux 1969; Taylor et al. 2001), *crispa* (*cri*; Lamm 1949; Tattersall et al. 2005), *tendril-less* (*tl*; Vilmorin 1910; Hofer et al. 2009), *cochleata* (*coch*; Wellensiek 1959; Couzigou et al. 2012), *apulvinic* (*apu*; Harvey 1979; Chen et al. 2012), *crispoid* (*crd*; Świecicki 1989; Berdnikov et al. 2000; McAdam et al. 2017) and *stipules reduced* (*st*; Pellew and Sverdrup 1923; Moreau et al. 2018). These diverse mutations affect various aspects of leaf form (Figure 1B-C): degree of pinnation (*uni* and *stp*), leaf polarity (*cri*), organ identity (*tl*, *apu, coch*) or organ patterning and development (*crd*, *st*).

**Figure 1.**
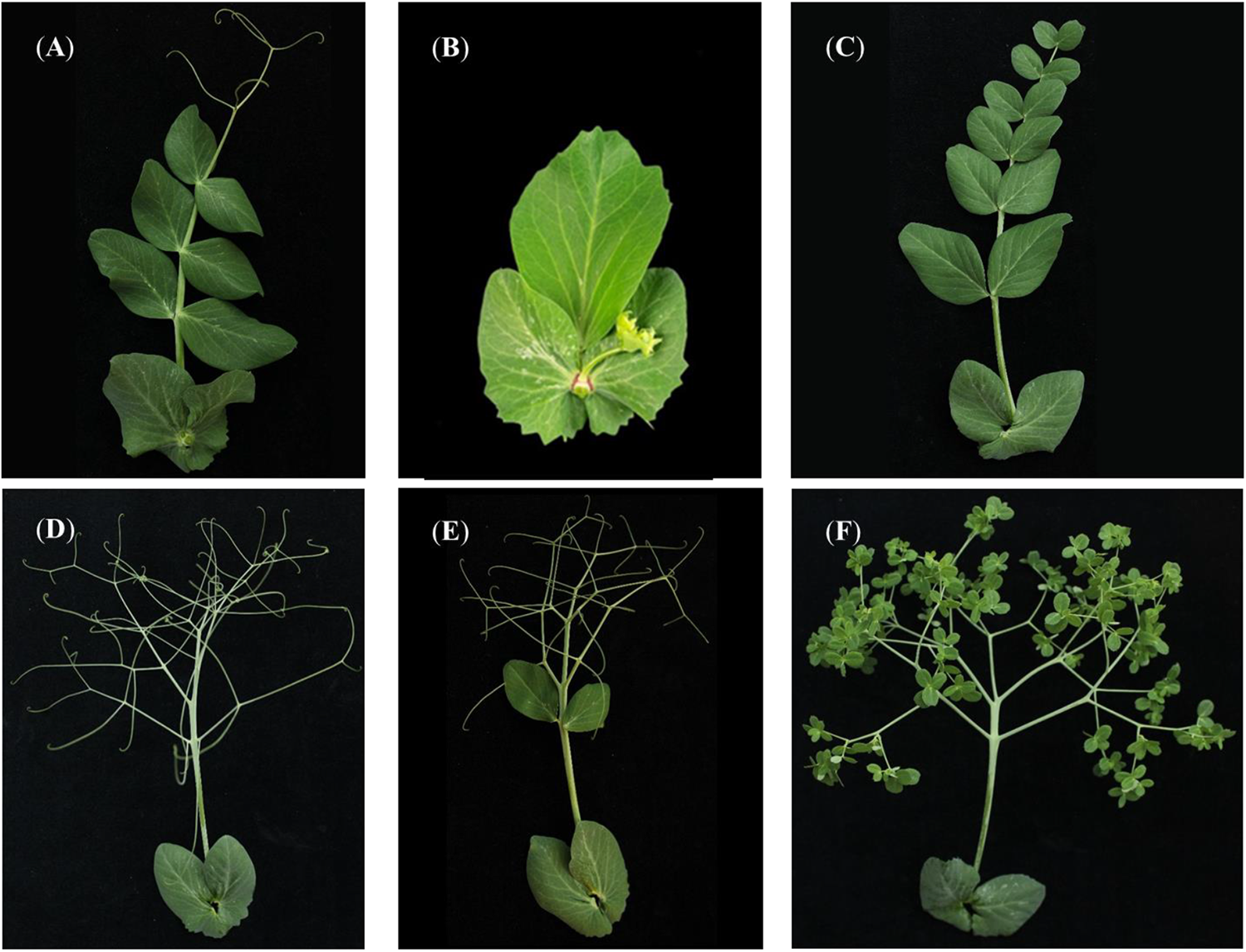
Mutants with altered compound leaf morphology in *Pisum sativum*. (A) A leaf of cv. Caméor representing conventional (wild type) pea leaf morphology. It is composed of two basal stipules, three proximal leaflet pairs, two distal tendril pairs and a terminal tendril. (B) A leaf from FN1210_1_42BC1 taken at a flowering node to show the combined effect of the *unifoliata* (*uni*) mutation on leaf and flower morphogenesis. The *uni* leaf has two basal stipules and a single leaflet. (C) A leaf from accession line VCC0350 carrying the *tendril-less* (*tl*) mutation. All tendrils are replaced by leaflets. (D) A leaf of cv. Kayanne exhibiting a semi-leafless phenotype resulting from the *afila* (*af*) mutation. No leaflets are present and tendrils are located on the branches (termed rachides or rachillae) of the leaf midrib. This phenotype is common to all currently available semi-leafless pea cultivars. (E) A leaf of accession JI 3129 that shows a pair of leaflets of normal size and morphology and the ramification of the central rachis. JI 3129, descending from cv. Raman, holds a novel *af* allele induced by a combination of gamma rays and fast neutrons (Ambrose 2004). (F) A leaf from accession NGB101745 carrying simultaneously *af* and *tl* mutations. It is representative of the pea ‘parsley leaf’ type. Seeds for accessions JI 2171 and JI 3129 were provided by the Germplasm Resources Unit at the John Innes Centre. Seeds for VCC0350 and NGB101745 were provided by INRAE CRB PROTEA, Dijon, France and Nordic Genetic Resource Center, Alnarp, Sweden, respectively. Photos are from this study.

One of the major successes of pea breeding in the past fifty years is attributed to the development of semi-leafless varieties carrying an *afila* (*af*) mutation, where leaflets are replaced by pinnae bearing tendrils (Snoad and Gent 1976; Kielpinski and Blixt 1982; Mikić et al. 2011; Tayeh et al. 2015a). Semi-leafless pea types are particularly suitable for large-scale production and mechanical harvesting; they exhibit a significantly increased standing ability because plants with strongly intertwined leaves can better support each other. They also show a reduced severity of foliar disease and enhanced uniformity of ripening and drying due to increased air movement and homogeneous light distribution inside the canopy (Snoad 1974; Mikić et al. 2011a). The yield superiority of semi-leafless types compared to conventional leafy types under normal competitive field conditions has been reported (Snoad 1974; Kielpinski and Blixt 1982), however, this type has been shown to be associated with reduced plant nitrogen content and seed size potential (Burstin et al. 2006). Contemporary dry pea breeding programmes worldwide almost exclusively use the semi-leafless ideotype. For instance, except for cultivar (cv.) Blueman, all dry pea cvs. registered in Canada (https://inspection.canada.ca/active/netapp/regvar/regvar_lookupe.aspx), France (https://www.geves.fr/catalogue-france/) and the UK (https://www.bspb.co.uk/wp-content/uploads/PGRO-Pulse-DL-Lists-2022.pdf) in the past five years are semi-leafless. On average from 1994 to 2021, Canada has been the world leader in dry pea production and France the third (FAOSTAT, 2022).

The semi-leafless (or afila) phenotype in pea arises as a consequence of a recessive mutation at a single locus (Kujala 1953; Solov’eva 1958; Goldenberg 1965; Khangildin 1966; Swiecicki 1982) on linkage group I (Marx 1969). In combination with *st* (Marx 1987), which reduces stipule size, the overall leaf lamina area becomes very reduced, and this is often (erroneously) referred to as a “leafless” phenotype; both *af* and *af,st* cvs. produce leaves at every node, but the morphology of these leaves is altered. In wild type (*Af*) genotypes, the proximally positioned lateral primordia, on the flanks of the leaf primordium, develop into determinate leaflet laminae, but in *af* mutants this fails to happen; the lateral and apical primordia behave in the same way such that second-order, midrib-like, primordia in turn initiate organ primordia (Meicenheimer et al. 1983; Côté et al. 1992). In a mature *af* leaf (that is otherwise wild type, e.g. carrying the dominant *Tl* allele), leaflets are replaced by pinnate structures bearing lateral and terminal tendrils, while the distal portion of the midrib that already bears lateral and terminal tendrils remains unchanged compared to a conventional leaf (Figure 1D).

There are three early reports of spontaneously occurring *af* mutants; the first was from Kujala (1953) based on a field observation in spring of 1950. The second report was from Solov’eva (1958) describing a ‘whiskered’ (усатый, usatyy, sometimes transliterated as usatyj) pea, first seen in 1949 (Kielpinski and Blixt 1982). The third report from Goldenberg (1965), described a phenotypically similar spontaneous mutation that occurred in cv. Cuarentona in Argentina. An early γ irradiation-induced *afila* allele was described by Jaranowski (1970) in the line Wasata-*af*.

*af* was mapped on pea linkage group (LG) I (Blixt 1972; Bordat et al. 2011; Tayeh et al. 2015b). The interaction of *Af* with other genes has been inspected in double or triple mutants (Marx 1974; Gourlay et al. 2000) (Figure 1F) in efforts to understand *Af* function. These suggested that *Af* is a negative regulator of *Uni* transcription (Hofer and Ellis 1998; Gourlay et al. 2000) and that *Af* is a negative regulator of auxin synthesis or implicated in auxin conjugation or catabolism (DeMason et al. 2013). Pinna initiation and determination during leaf development in pea and the establishment of pinna identity are thus likely to depend on auxin gradients together with the antagonistic interaction between *Uni* and *Af* (Hofer and Ellis 1998; Gourlay et al. 2000; DeMason et al. 2013). In *M. truncatula*, a close relative of pea, it has been shown that a Cys(2)His(2)-zinc finger transcription factor named PALM1 binds directly to the promoter of *SINGLE LEAFLET1* (*SGL1*), the orthologue of *Uni*, thereby repressing *SGL1* expression (Chen et al. 2010; He et al. 2020).

Besides the lack of information on the molecular identity of *af*, it was also unclear how the pea community managed to transform *af* from simple spontaneous mutations observed in the USSR (Solov’eva 1958), Finland (Kujala 1953) and Argentina (Goldenberg 1965) into a major breeding criterion in dry pea. Did pioneer breeders simply rely on spontaneous mutations in their programmes? What part did mutation breeding (Blixt 1972) play in developing semi-leafless pea varieties? Multiple induced mutants from different genetic backgrounds exhibit phenotypes similar to the spontaneous *af* mutants (Figure 1E; Kielpinski and Blixt 1982; Ambrose 2004) and are available in germplasm collections. Therefore, the aim of this study was to: (1) shed light on the genes underlying the afila phenotype and thus contribute to the identification and description of one of the key elements of the determination of leaf formation in pea; and (2) highlight the origin and prevalence of various alleles and haplotypes associated with the semi-leafless phenotype in pea crop breeding.

## Materials and Methods

### Plant material

The pea cvs. Ballet, Terese, and Baccara show a semi-leafless phenotype while cvs./landraces Cameor, Champagne, PI180693 and VavD265 have a conventional leaf type. Four RIL populations, namely Pop2, Pop4, Pop5, and BxP, obtained by single seed descent from crosses between these accessions (Tayeh et al. 2015b), were used in this study (Supplementary Table 1). The 450 RILs from all 4 populations were evaluated for leaf type and phenotypic data were used to place *af* on the consensus map from Tayeh et al. (2015b). In an independent experiment, a 58-individual RIL population, derived from the cross between JI1201 and JI813 from the John Innes *Pisum* germplasm, was used for the same purpose together with 967 F_7_ and F_8_ individuals, segregating for *Af/af* obtained by selfing a single F_4_ heterozygote line (Supplementary Table 1; Data S1).

Fifty-nine accessions with conventional leaf type and 107 accessions carrying the *af* mutation, including 63 and 72 % cvs., respectively, were used as a validation panel to confirm the identity of the candidate molecular determinants for the *af*-related phenotype and to assess the different haplotypes at this locus (Supplementary Table 4; Fig. 4). Cvs. were bred in Europe, North America and Australia. Different seed sources were taken for some mutants and cvs. for confirmation and thus the total number of examined accessions is 173 (Supplementary Table 4). Seeds were provided by CRB PROTEA at INRAE (France), the John Innes centre (United Kingdom) and the Nordic Genetic Resource centre (Sweden).

EMS mutant alleles of *Af* candidate genes, *PsPALM1a* and *PsPALM1b*, in cv. Cameor were identified using a TILLING approach. M3 mutant lines were screened for substitutions in the coding sequences of these genes and homozygotes were selected in the M3 or M4 generation. All recovered homozygous plants, 51 for *PsPALM1a* and 37 for *PsPALM1b*, were subjected to leaf phenotyping in a glasshouse. Three lines carrying mutations provoking non-synonymous amino acid changes or premature stop codons were selected for further analyses. Two cycles of backcrossing to cv. Cameor were performed. Homozygotes for the mutant alleles were then crossed to a semi-leafless accession (JI1195) where both *PsPALM1a* and *PsPALM1b* are lacking. The F_1_ plants, hemizygous for either one wild type *PsPALM1b* allele and one mutant *PsPALM1a* allele, or, one wild type *PsPALM1a* allele and one mutant *PsPALM1b* allele, were analysed for their leaf types. Gamma-induced mutants from cvs. Auralia, Borek and Raman and line FN2070, a fast neutron-induced palmate mutant derived from accession JI2822, were also used for phenotyping and molecular analyses. Gamma-ray induced mutants were provided by the Bulgarian Academy of Sciences, Ulpia Serdica, Sofia-Capital, Bulgaria and the fast neutron mutant resource has been described by Domoney et al. (2013).

### DNA extraction and SNP genotyping

DNA extraction from fresh young leaves, collected from the different accessions in the plant material section, was conducted using Macherey-Nagel™ NucleoSpin™ Plant II mini kit (Macherey-Nagel, Düren, Germany) following the manufacturer’s instructions. Elution was performed in 50 μL ultrapure water. DNA purity and concentration were evaluated using a NanoDrop™ ND-1000 spectrophotometer (Thermo Scientific, Wilmington, United States).

Polymerase chain reactions (PCR) were conducted to determine the presence or absence of seven candidate genes located in the *af*-harbouring region. Their products were run on agarose gel for visual analysis. PCR Primers were designed based on the pea transcript sequences from Alves-Carvalho et al. (2015) that share the highest homology with the sequences from model legumes *M. truncatula* and *G. max* (see ‘Synteny-based ortholog identification’ section). A PCR primer set related to a sucrose phosphate synthase (SPS) gene located outside the *af* region and on a separate chromosome (Aubert et al. 2006) was regularly used as a control to rule out any problems in the presence/absence detection related to DNA or PCR mix. Primer sequences, corresponding genes, and expected product sizes are available in Supplementary Table 3. The SPS marker located on LG IV (Aubert et al. 2006), was used as a PCR control when checking the presence/absence of genes from the *af* locus.

### cDNA-AFLP mapping

RNA was extracted from apical and lateral shoot tips of approx. 5 mm in length harvested from 1 month old JI 1201 x JI 813 RIL seedlings, 2 weeks after decapitation. Double-stranded cDNA was synthesised from 2 μg of total RNA using AccuScript High Fidelity 1st Strand cDNA Synthesis Kit (Stratagene, La Jolla, USA) according to manufacturer’s instructions, except that RNase inhibitor was omitted and reverse transcription was carried out using a polyA-specific primer containing a PstI site as described in Money et al. (1996). AFLP template preparation was performed as described by Vos et al. (1995) using cDNA digested with *Pst*I and *Mse*I restriction enzymes (Boehringer, Mannheim, Germany), followed by adapter ligation and amplification with 2 bases of selection on the γ33P-ATP-labelled *Pst*I primer and 2 bases of selection on the *Mse*I primer. AFLP bands were separated on a polyacrylamide gel according to Bachem et al. (1996).

### Synteny-based candidate gene identification

The transcript sequences corresponding to the *af*-flanking SNP markers from the consensus map of Tayeh et al. (2015b) were used for ortholog search in *M. truncatula* Mt4.0 genome release (Tang et al. 2014) and *G. max* Williams 82 assembly 2 accessible at SoyBase (Grant et al. 2010). In all cases, best hits from reciprocal BLAST were considered as orthologs. Gene order on *M. truncatula* and *G. max* pseudomolecules was cross-compared with the gene order on *P. sativum* v1.0 genome assembly version (Kreplak et al. 2019). To assess the extent of synteny conservation, *Arabidopsis thaliana* genome assembly available from The Arabidopsis Information Resource (Berardini et al. 2015) was further included for ortholog search. *Lens culinaris* CDC Redberry Genome v1.2 available on the KnowPulse web portal (http://knowpulse.usask.ca) and *Cicer arietinum* Desi uwa-V3.0 genome assembly version (Edwards 2016) were also used for comparative genomics.

### Fast neutron mutant analysis

Raw fastq reads of the FN2070 palmate mutant (ENA accession reference PRJEB60035) were mapped using bwa mem (v 7.0.12; Li and Durbin 2010), and the bam file was processed using samtools (v1.9; Danecek et al. 2021). The pea ZW6 genome (Yang et al. 2022) was split into 100kb windows using bedtools (v2.2.28; Quinlan and Hall 2010), and samtools cov was used to find the coverage in these windows. The deletion on LG I, at Chr2:473791227-474324688, was curated manually from the raw data.

### Virus-induced gene silencing of the two *PsPALM1* candidate genes

A 349-bp DNA fragment from the pea cv. Champagne *PsPALM1a* gene was PCR-amplified using primers PALM1aF2R2-fwd (5’-cgggatccCATCATCCTTCTTCTCCATTC-3’) and PALM1aF2R2-rev (5’ cgggatccCCAAACGTAGCTCAAGATCG 3’). The 349-bp fragment of *PsPALM1a* shares 95.4% nucleotide identity with *PsPALM1b* and is located at the 3’ end of the two genes. The PCR product was cloned into pGEM-T plasmid (Promega) and sequenced. This 349-bp fragment was then subcloned into the *BamH*I cloning site of the pBPMV-IA-V1 plasmid (Zhang et al. 2010). Orientation of the insert was determined by PCR using a combination of vector-specific (RNA2-BamH1-rev: 5’ AGCATACTCAACGAG AGGGTCA 3’) and fragment-specific (PALM1aF2R2-rev) primers. The insert sequence and its reverse orientation was confirmed by Sanger sequencing (GATC, Germany).

BPMV-*PsPALM1* VIGS vector was delivered into *Phaseolus vulgaris* cv. Black Valentine plants by mechanical inoculation of the two BPMV-derived infectious plasmids: pBPMV-IA-R1M (carrying RNA1 of BPMV) and pBPMV-*PsPALM1* (carrying RNA2 of BPMV with the inserted 349bp-*PsPALM1a* fragment) (Pflieger et al. 2014, 2015). After 3 weeks, infected *P. vulgaris* leaves were used as viral inoculum to inoculate 15 healthy pea cv. Champagne plants as described previously (Meziadi et al. 2016, 2017). Nine mock-inoculated (Mock) and nine wild type BPMV inoculated (BPMV-0) plants were used as controls. All plants were pruned at 4 weeks post-inoculation in order to concentrate BPMV titre and increase VIGS efficiency (Meziadi et al. 2016, 2017). Leaf morphology was scored at 11.5 weeks post-inoculation. A Chi^2^ statistic test was used for data analysis.

### Quantitative real-time PCR

Quantitative real-time PCR were run on Biomark™ HD system (Fluidigm) according to manufacturer’s protocol to assess the relative gene expression of *PsPALM1a*, *PsPALM1b* and *Uni.* Total RNA extraction from different tissue samples collected from pea cv. Cameor (D’Erfurth et al. 2012; Alves-Carvalho et al. 2015; Vernoud et al. 2021) was performed using the same protocol from Gallardo et al. (2007). Tissue samples included: shoots, roots, nodules, developing seeds (8-29 days after pollination (DAP)), dissected seeds at 12 DAP (embryo, albumen, tegument) and seeds at 3 days post-imbibition. Three biological replicates were included. Relative expression levels were evaluated using 2^−ΔΔCt^ method and three control genes, i.e. *histone H1*, *actin* and *EF1α*, were used for data normalization as described by Alves-Carvalho et al. (2015). For investigating gene expression in EMS TILLING mutant 2835 and Bulgarian gamma-ray mutants in cvs. Auralia, Borek, and Raman, ten germinating seeds at three days post-imbibition were considered per condition with each being a separate biological replicate.

## Results

### *af* is located within a 1.4-cM interval on pea linkage group I with PsCam023334_13180_1532 as the closest molecular marker

Five pea recombinant inbred line (RIL) populations (Ellis et al. 1992; Tayeh et al. 2015b) (Pop2, Pop4, Pop5, BxP and JI1201xJI0813) segregate alleles at the *af* locus (Supplementary Table 1). One parent from each population has a conventional leaf type as in Figure 1A and the other exhibits a semi-leafless phenotype as in Figure 1D or the *af, tl, s*t triple mutant in the case of JI1201. The phenotyping and genotyping data obtained for Pop2, Pop4, Pop5 and BxP placed the *af* mutation at 85.6 cM on linkage group I (Chromosome 2) of the integrated pea consensus genetic map from Tayeh et al. (2015b) (Figure 2A). Thirteen out of the 450 total F_6-9_ inbred lines carry recombination events near the morphological marker *af* (Figure 2B) and were further examined. Based on their haplotypes inferred from markers exclusively polymorphic in all four populations including *af* (Figure 2C), the *Af*-containing region was narrowed down to a 1.4-cM interval located between gene-based SNP markers PsCam023334_13180_1532 and PsCam000001_1_322.

**Supplementary Table 1.** Recombinant inbred line populations derived by single seed descent from crosses between parents contrasting for their leaf types. The parent indicated in green carries an *af* allele. The other parent has a conventional leaf type (*Af*).

| Population ID | Female parent | Male parent | Generation | Number of individuals | References |
| --- | --- | --- | --- | --- | --- |
| Pop2 | Champagne | Terese (*af*) | F8:9 | 76 | Loridon et al. 2005; Tayeh et al. 2015b |
| Pop4 | Ballet (*af*) | Cameor | F6:8 | 159 | Bourion et al. 2010;<br>Tayeh et al. 2015b |
| Pop5 | VavD265 | Ballet (*af*) | F6:8 & F6:9 | 168 | Bordat et al. 2011;<br>Tayeh et al. 2015b |
| BxP | Baccara (*af*) | PI180693 | F8 | 47 | Hamon et al. 2011;<br>Tayeh et al. 2015b |
| Jl1201XJl0813 | Jl1201<br>(*af,tl, st*) | Jl0813 | F4 & F8 | 58 & 967 | Ellis et al. 1992;<br>This study |

**Figure 2.**
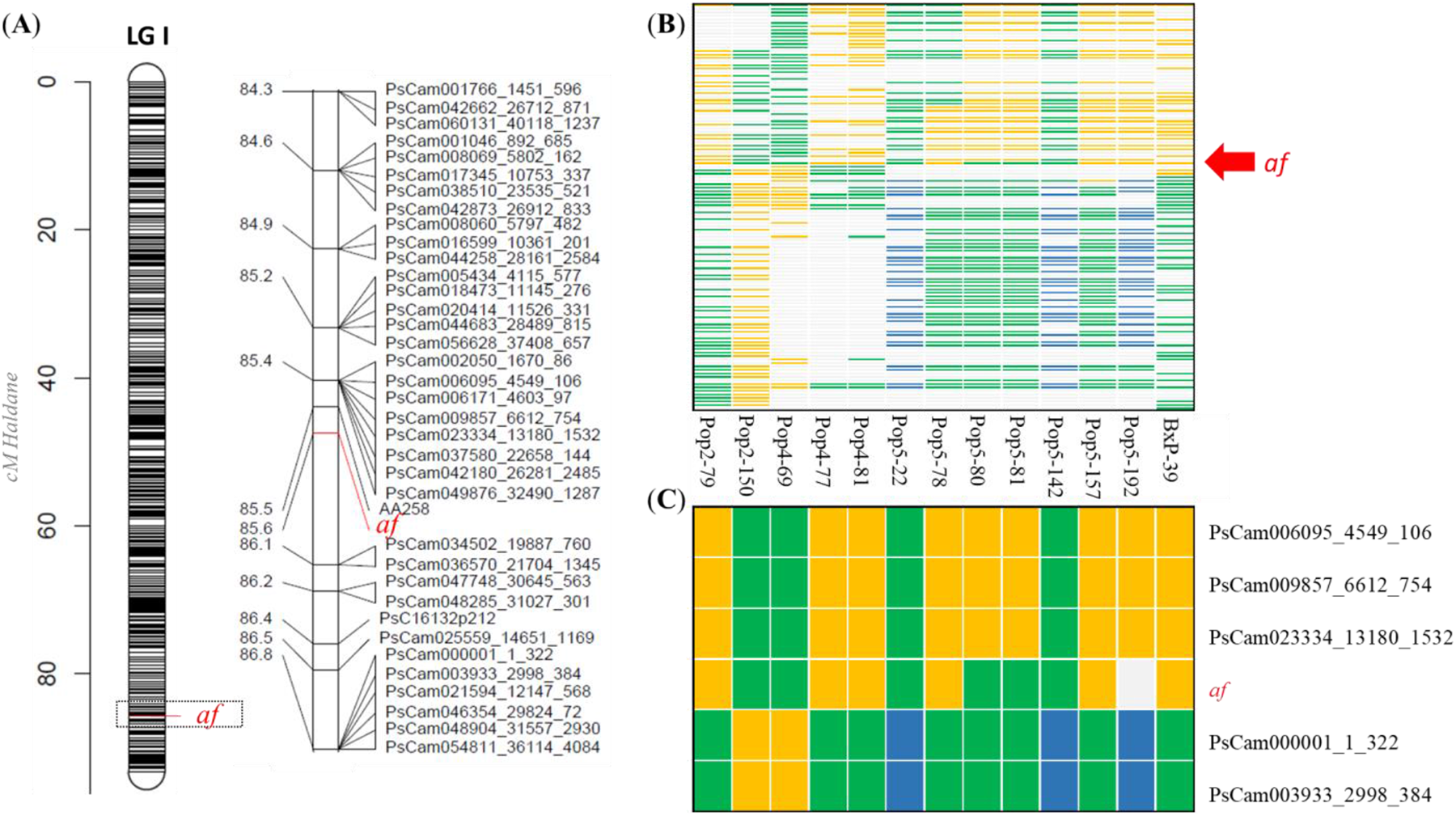
Fine mapping of the *af* locus. (A) Schematic representation of marker density on pea linkage group I on the integrated consensus map of Tayeh et al. (2015b); the region near *af* is shown in detail. Maps were constructed using LinkageMapView R package (Ouellette et al., 2018). (B) Graphical genotype illustration for 13 recombinant inbred lines carrying recombination events near the *af* locus. Lines are from four different populations: Pop2, Pop4, Pop5 and BxP. The orange colour refers to alleles inherited from parents characterized by a conventional leaf type. The green colour refers to alleles inherited from parents showing semi-leafless phenotypes. The blue colour indicates heterozygosity. The gray colour is used when data are not available: monomorphic markers or missing data. (C) Graphical genotype illustration considering only the closest markers to *af* and polymorphic in all four mapping populations. The order of the individuals is the same as in section B. The position of *af* in sections A-C is marked in red.

*af* is genetically closer to PsCam023334_13180_1532 than to PsCam000001_1_322: only 2 recombination events were detected between *af* and PsCam023334_13180_1532 whereas 10 recombination events were detected between *af* and PsCam000001_1_322. Because of the recessive nature of the *af* mutation and residual heterozygosity, the exact position of the recombination event in the thirteenth line with recombination near *af*, viz. Pop5-192, could not be determined (Figure 2C). In the 58 F_4_ RILs derived from the cross between JI1201 and JI0813, cDNA-AFLP markers corresponding to transcripts PsCam048460 and PsCam042853 (Alves-Carvalho et al. 2015) were mapped close to *af*. A fine mapping population of 967 F_7_ individuals was derived from an F_4_ *Afaf* heterozygous plant (Supplementary File 1). No recombinant was detected between a marker corresponding to PsCam002050 and *af*. Interestingly, a SNP marker located in PsCam002050, namely PsCam002050_1670_86, co-locates with PsCam023334_13180_1532 on the pea consensus map (Tayeh et al. 2015b).

### Co-orthologs of M. truncatula PALMATE-LIKE PENTAFOLIATA1 (PALM1) are candidate genes for Af

Candidate genes for *Af* were mined using *af* flanking markers from the pea consensus genetic map and the synteny relationships with close species with available genome assemblies (Figure 3; Supplementary Table 2). *Medicago truncatula* and *Glycine max* were considered. Fifty-two and 32 genes were identified in the synteny blocks on *M. truncatula* chromosome 5 and *G. max* chromosome 11, respectively (Supplementary Table 2), including orthologues of the PsCam002050-related gene, *Medtr5g014260* and *Glyma11g04530*. As highlighted in Figure 3, collinearity was overall conserved between pea, *M. truncatula*, *G. max* and even *Arabidopsis thaliana*. Medtr5g014400, named *PALMATE-LIKE PENTAFOLIATA1* (*PALM1*), was one of the genes located in the collinear block from *M. truncatula*. It is the equivalent of MtrunA17_Chr5g0400571 in the latest r5.0 assembly version of the *M. truncatula* A17 genome and has conserved orthologs in *G. max* and *A. thaliana* (Figure 3). Two strong lines of evidence indicated that the pea ortholog of *Medtr5g014400* is a relevant candidate gene for *Af*: 1) *Medtr5g014400* was reported to control compound leaf morphology in *M. truncatula* (Chen et al. 2010), and 2) *Medtr5g014400* is only 68.9 and 15.5-kb distant from *Medtr5g014260* and *Medtr5g014410*, the orthologues of the pea genes corresponding to *af* flanking markers, of which, PsCam023334_13180_1532 is the closest to *af* (Supplementary Table 2). Contrary to *M. truncatula, G. max* and *A. thaliana*, not one but two sequences similar to *PALM1* were identified by homology search against pea genomic sequence data from cv. Cameor (Aluome et al. 2016). The putative genes were named *PsPALM1a* (scaffold_45110) and *PsPALM1b* (scaffold_30425). Three copies of the neighbouring gene, encoding an N2-acetylornithine deacetylase, were identified in pea, whereas *M. truncatula* and *G. max* each have only a single copy (*Medtr5g014410* and *Glyma11g04650*). Pea genes predicted to encode N2-acetylornithine deacetylase according to the PsCam_LowCopy Unigene set (Alves-Carvalho et al. 2015) are hereafter referred to as: *PsNAOD1* (PsCam045967), *PsNAOD2* (PsCam027203) and *PsNAOD3* (PsCam023334).

**Supplementary Table 2.** See separate file. List of gene-based SNP markers tightly-linked to *af* and predicted genes in collinear regions in *M. truncatula*, *G. max* and *A. thaliana*. Each row contains a distinct ortholog set. When only one gene ID is indicated in a complete row, this means that no orthologs could be identified from other species. The assembly versions used to construct this table are as follows: Mt4.0 for *M. truncatula*, Williams 82 Assembly 1 for *G. max* and TAIR10 for *A. thaliana*.

**Figure 3.**
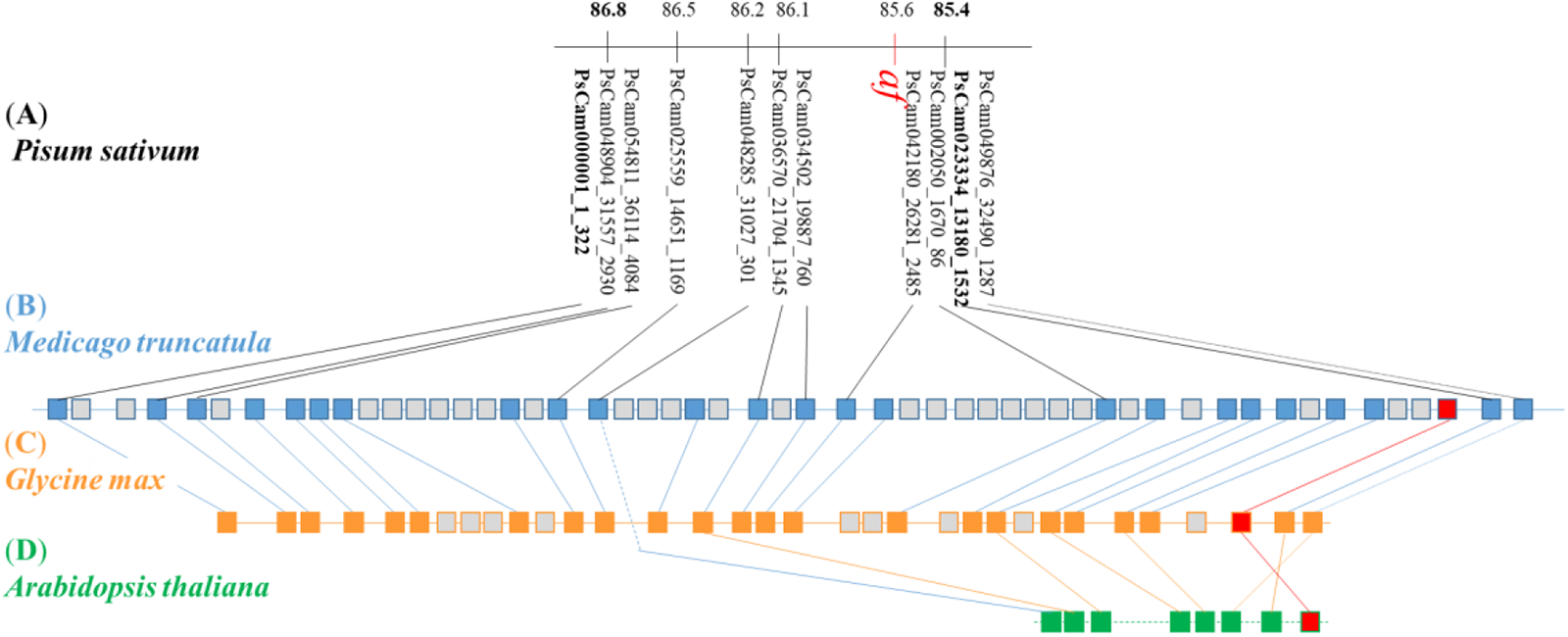
Collinearity between the pea linkage map at the *af* locus and the physical maps of *M. truncatula* chromosome 5, *G. max* chromosome 11 and *A. thaliana* chromosome 4. (A) Order of the gene-based SNP markers on the integrated consensus genetic map at the *af* locus. The polymorphic markers in the four RIL populations Pop2, Pop4, Pop5 and BxP, delimiting the fine-mapped *af* interval, are in bold. (B-D) Schematic illustration of the order of genes in the syntenic regions in *M. truncatula* (B), *G. max* (C) and A*. thaliana* (D). These regions are of total length of 407, 331 and 101 kb, respectively. Colour-filled boxes refer to predicted genes that have ortholog counterparts in the studied species. Black, blue and orange lines are used to link putative orthologs. Grey boxes refer to predicted genes that do not have equivalents in other species. *A. thaliana* is used as an outgroup to show the extent of collinearity conservation beyond the legume family. In total, 30 genes are contained in the illustrated genomic region from *A. thaliana* chromosome 4 but, for the sake of clarity, only those having putative orthologs in *M. truncatula* or *G. max* are illustrated. The orthologs of *PsPALM1a* and *PsPALM1b*, candidate genes for *Af*, are shown as red squares.

### *PsPALM1a*, *PsPALM1b* and *PsNAOD1* are absent in semi-leafless parent and recombinant inbred lines

Primer pairs were designed to clone *PsPALM1a* and *PsPALM1b* from parental lines of the aforementioned RIL populations (Supplementary Table 3). The goal was to identify potential causal mutations in the coding and regulatory regions that could possibly explain the *af* leaf phenotype. Interestingly, no amplicons could be obtained from semi-leafless lines, whereas amplicons had the expected sizes from accessions with conventional leaves (Supplementary Figure 1A). The same observation was made with primers for *PsNAOD1* (Supplementary Figure 1B). Genotyping the recombinant lines from Pop2, Pop4, Pop5 and BxP at the *af* locus (Figure 2C), taking advantage of the presence/absence polymorphism for these three genes, confirmed the co-segregation of *PsPALM1a*, *PsPALM1b* and *PsNAOD1* with *Af*. In fact, all semi-leafless RILs showed absence of amplification. RILs with conventional compound leaves, taken as control, showed normal amplification. Furthermore, mapping whole-genome sequence reads from pea accessions included in the SNP detection panel of Tayeh et al. (2015b) against cv. Cameor reference genomic scaffolds revealed an absence of read coverage for the semi-leafless genotypes for the scaffolds corresponding to *PsPALM1a, PsPALM1b,* and *PsNAOD1* whereas read coverage was normal for all conventional-leaved genotypes (Supplementary Figure 1C). The results thus suggested that the deletion of at least three genes, including the co-orthologs of *M. truncatula PALM1*, is tightly linked to the *af* mutation in pea.

**Figure S1.**
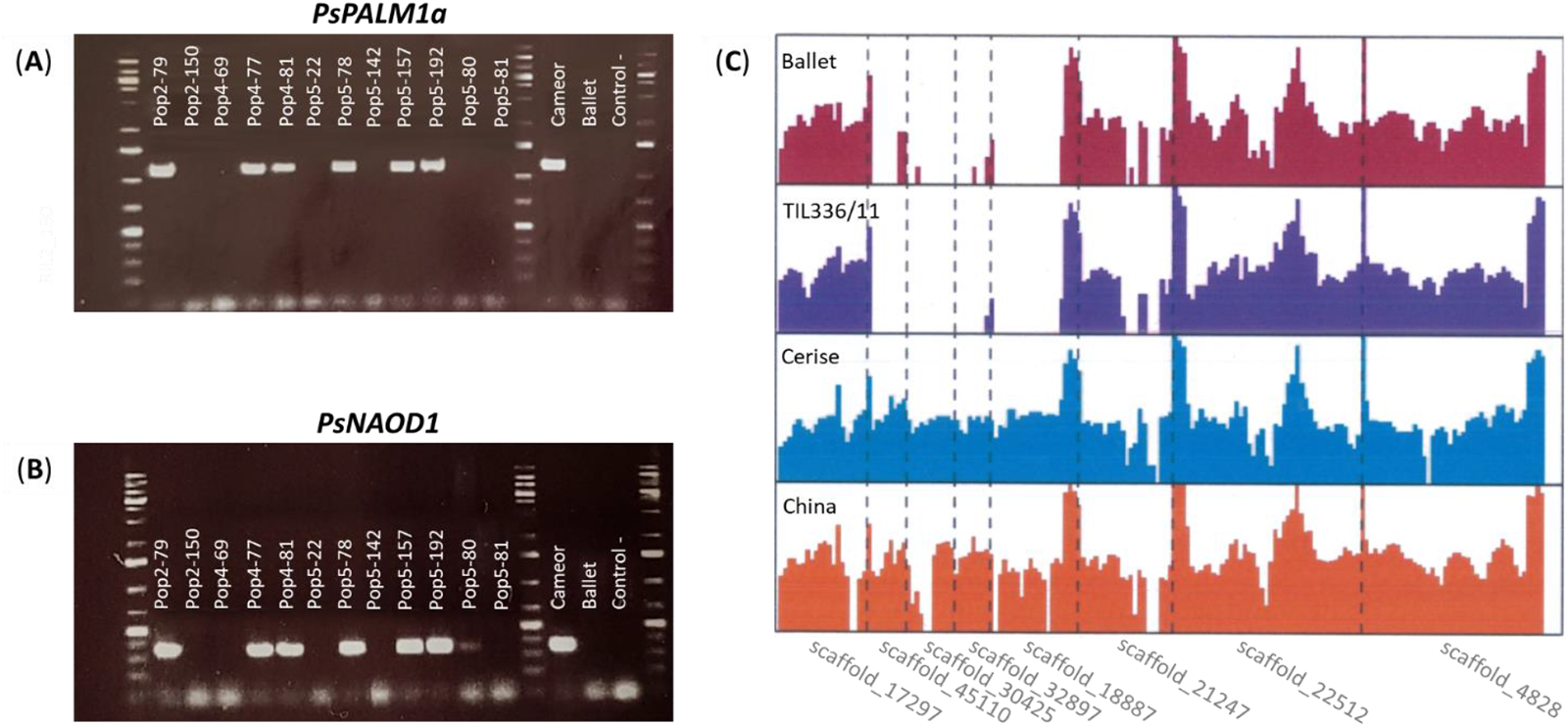
Presence versus absence of *PsPALM1a, PsPALM1b* and *PsNAOD1* in conventional and semi-leafless pea accessions. (A-B) Agarose gel electrophoresis of PCR products amplified using specific primers for *PsPALM1a* (A) and *PsNAOD1* (B). The size of the obtained products is measured with DNA ladders in lanes 1, 14 and 18. Lanes 2 to 13 show PCR products from recombinant inbred lines with conventional or semi-leafless phenotype. Lines with conventional leaf type are Pop2-79, Pop4-77, Pop4-81, Pop5-78, Pop5-157 and Pop5-192. Cameor (conventional) and Ballet (semi-leafless), in lanes 15 and 16, were used as positive and negative controls, respectively. Lane 17, without DNA template, represents an additional negative control. (C) Mapping of genomic reads obtained through whole-genome sequencing of cultivar Ballet and breeding lines TIL336/11, Cerise and China against genespace scaffolds (Aluome et al. 2016). The latter two accessions have a conventional leaf type whereas the former two are semi-leafless. Scaffold_45110, scaffold_30425 and scaffold_1887 harbour *PsPALM1a*, *PsPALM1b* and *PsNAOD1*, respectively. The other scaffolds include other genes expected to be near *Af* based on synteny analysis.

**Supplementary Table 3.** See separate file. Primers used to examine the presence/absence variations at the *af* locus and to quantify gene expression. Primer pairs #1 to 8 were used to run standard amplifications on genomic DNA. The order of primer pairs #1 to 7 follows the expected order of corresponding genes on pea Chromosome 2. Primer pair #8 amplifies a control gene that was selected out of chromosome 2. Primer pairs #9 to 11 were used for gene expression detection and quantification. Primer pairs #12 to 13 were used for fine mapping as reported in Data S1.

### Eight haplotypes define the genomic region harbouring the *af* mutation and show that deletion of *PsPALM1a* and *PsPALM1b* confers the af phenotype

To further extend the results obtained above, the presence/absence of seven consecutive pea genes, including *PsPALM1a* and *PsPALM1b*, was assessed on 166 accessions with conventional or leafless/semi-leafless phenotypes (Supplementary Table 4). These accessions included cvs., breeding lines and mutants of different geographical origins. The SPS marker located on LG IV (Aubert et al. 2006), was used as a positive PCR control. In total, eight haplotypes were identified in this study, with none to seven genes being absent (Figure 4). All genes examined were present in all the accessions presenting a conventional leaf type. These latter were thus assigned to a single haplotype, referred to as Haplotype 1 (Figure 4). *PsPALM1b* was the only gene to be systematically absent in all leafless or semi-leafless accessions (Haplotypes 2-8). *PsPALM1a* was similarly absent in all *af* cvs. but was present in three Bulgarian gamma ray-induced mutant lines derived from cvs. Auralia (Haplotype 6), Borek and Raman (Haplotype 7). The mutation in cv. Raman was confirmed by Ambrose (2004) as a new, weak allele at the *af* locus. The mutants derived from cvs. Auralia and Borek were not investigated in detail but occasionally their compound leaves produced a leaflet (Supplementary Figure 2).

**Figure 4.**
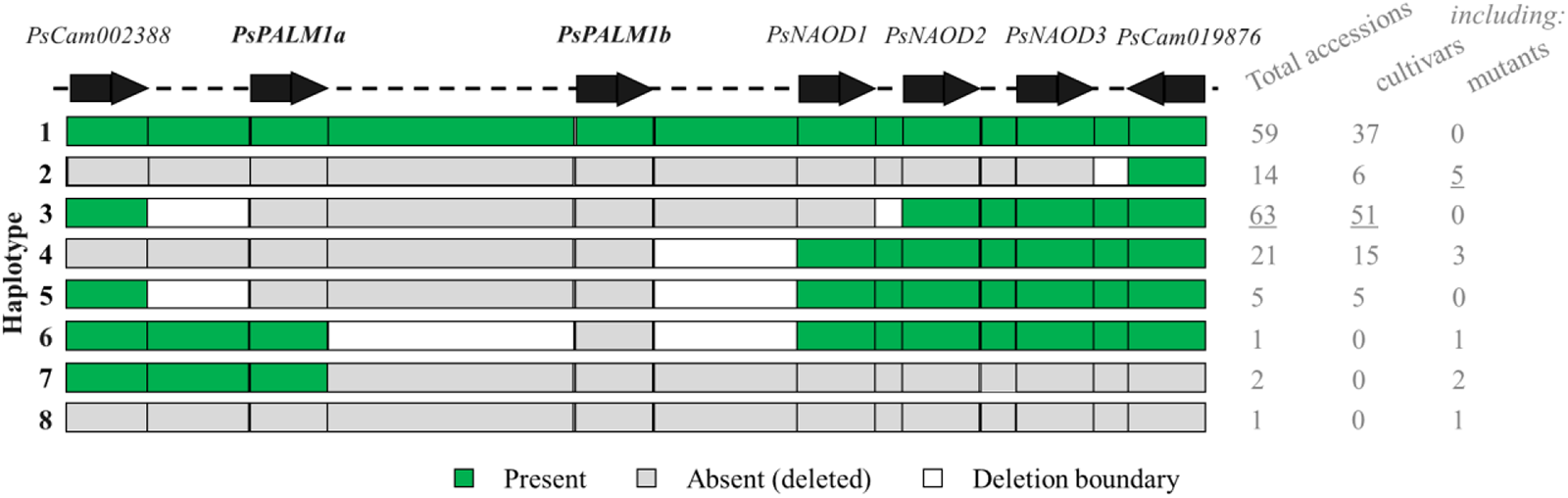
Diversity of haplotypes at the *af* locus. Black arrows represent *Af* candidate genes (*PsPALM1a* and *PsPALM1b*) and their neighbouring genes identified following a synteny-based strategy. Haplotypes are numbered 1 to 8. Green and grey boxes indicate whether the corresponding genes/regions are present or absent, respectively. White boxes represent intergenic regions where the deletion endpoints lie, but the positions are uncertain. It is noteworthy that the haplotypes in this figure are based on information obtained from PCR with specific primers for the seven examined genes. Each haplotype may have several subgroups once the deletion boundaries are precisely defined. All accessions with conventional leaf type are clustered under haplotype 1. Haplotypes 2-8 are exclusively found in leafless or semi-leafless accessions. The number of cultivars and mutants belonging to each haplotype, as well as the total number of examined accessions, are provided at the right side of the figure (see Supplementary Table 4).

**Supplementary Table 4.** See separate file. List of accessions examined for their haplotypes at the *af* locus. Conventional (*AFAF StSt*), semi-leafless (*afaf StSt*) or leafless (*afaf stst*) wild, landrace, germplasm, breeding and cultivar accessions are included. Each haplotype is distinguished by a separate color. When applicable, rows corresponding to the same accessions but where seeds were provided by different centres and used to confirm particular findings are colored differently. Lineages of pea accessions in the comment section are obtained from the donors, from Leterme et al. (1992; Anim. Feed Sci. Technol. 37:309-315) or from Cupic et al. (2009; Journal of Food, Agriculture & Environment 7:343-348). Each number under the gene name in columns 17 to 24 corresponds to the product size in bp obtained by PCR for each gene (primer pair) and each accession. It is noteworthy that the number “0” indicates no amplification and suggests that the gene is deleted. CRB PROTEA refers to the Pulse genetic resource unit at INRAE. JIC refers to the Germplasm Resources Unit at the John Innes Centre. NGB stands for Nordic Gene Bank and refers to the Nordic Genetic Resource Centre.

**Figure S2.**
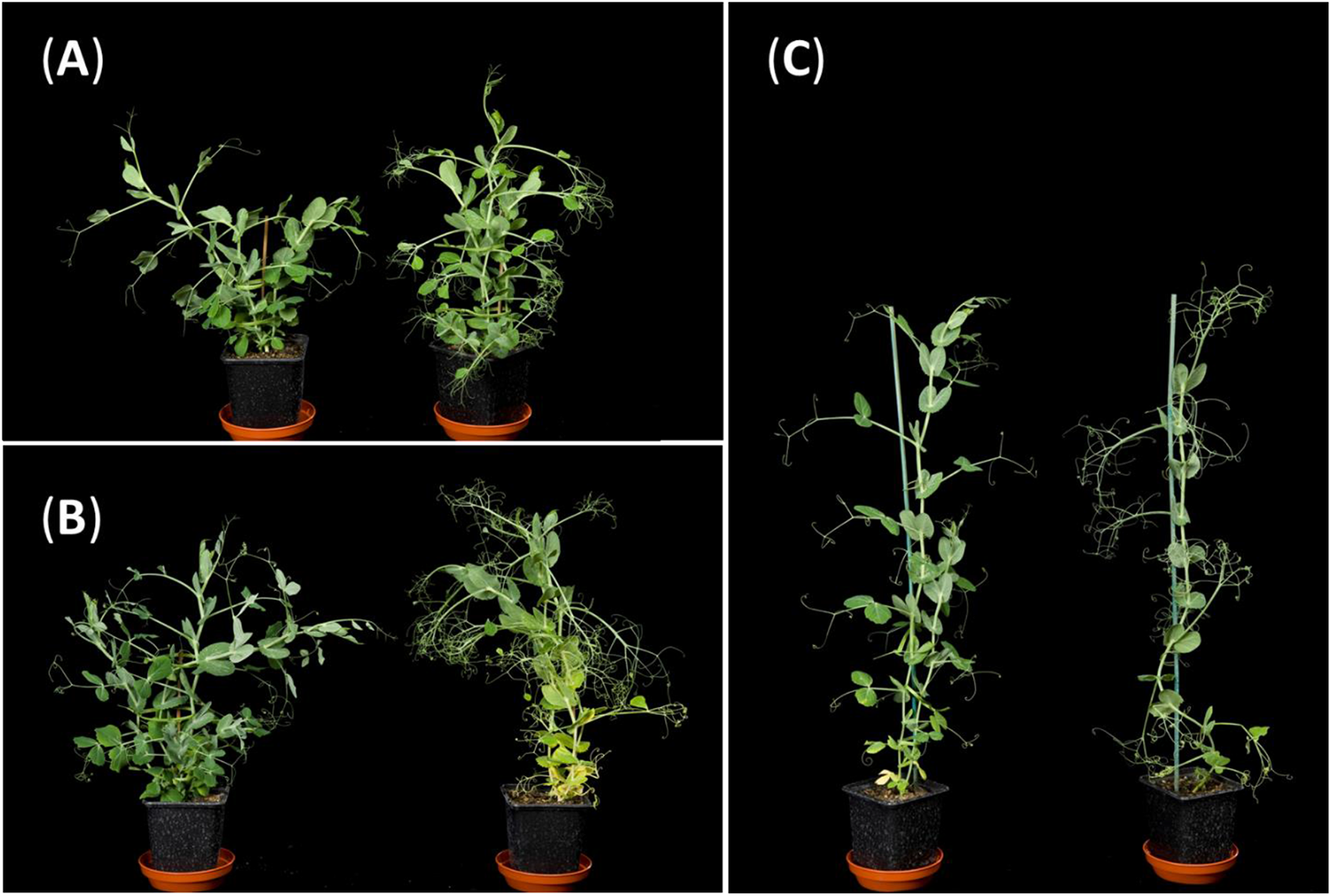
Gamma-ray-induced *afila* mutant lines originating from Bulgaria, carrying a deletion of *PsPALM1b*, shown in comparison to their corresponding wild type progenitor cultivars. (A) JI 3133, also known as cv. Raman (left); JI 3129, also known as line 11/47, with characteristic weak *af* leaf phenotype, where the leaf rachis usually bears proximal leaflets (right). (B) JI 3520, also known as cv. Borek (left); JI 3509, also known as line 2/544, with *af* phenotype and very occasionally additional proximal or distal leaflets (right). (C) JI 3132, also known as cv. Auralia (left); JI 3504, also known as line 1/534, with *af* phenotype (right).

For three haplotypes, the origins of the *af* alleles can be identified as follows:

1. Four accessions designated Usatuj or Usatyj are in haplotype 2; presumably this is Solov’eva’s Usatyj-5 (Solov’eva 1958). We propose to refer to this spontaneous mutant allele as *af^S^*. Haplotype 2 is carried by six registered cvs.
2. Haplotype 3 includes 51 cvs., of which Filby, Barton and Melton are recorded as being derived from a cross with an ‘Argentinian af’ obtained from G.A. Marx at Cornell (Supplementary Table 4). This haplotype also includes Marx’s isogenic series MISOG-1 and MISOG-2. This is consistent with Marx having obtained the *af* mutants from Goldenberg (Marx 1969); we can designate this allele as *af^G^*.
3. Haplotype 4 includes 15 cvs. and the line Wasata-af; this is clearly Jaranowski’s induced mutant (Jaranowski 1970) and can be designated *af^J^*.

Haplotype 5 is represented by five cvs. and must have an origin distinct from haplotypes 2, 3 or 4. The origin of this *af* allele is unknown. The *af* allele described by Kujala (1953) has no known descendants. Kujala’s 1953 paper suggests that this mutant was lost, so it would be unwise to suggest that this haplotype is connected to it.

Haplotypes 6, 7 and 8 occur in more recently generated mutant lines.

We note that the categorisation of some cvs. and germplasm lines is not consistent with their proposed origins (Supplementary Table 4). In breeding programmes involving *af* mutants from different sources, the alleles would have been phenotypically indistinguishable which may have led to these inconsistencies.

In all cases, these results are consistent with the af phenotype being the consequence of the loss of both co-orthologs of *M. truncatula PALM1*, i.e. *PsPALM1a* and *PsPALM1b*, and that different sources of *af* mutations have been used for breeding programs in the past half-century.

### *PsPALM1a* and *PsPALM1b* encode highly similar Q-type Cys(2)His(2)-zinc finger proteins with conserved functional domains and putative auxin response elements in their promoter regions

*PsPALM1a* and *PsPALM1b* are intron-less genes, predicted to encode 242 and 239-amino-acid proteins from 729 and 720-bp coding sequences, respectively. In cv. Cameor, they share 93.3 % identity at the nucleotide level and 90.7 % at the protein level. PsPALM1a and PsPALM1b each contain a single, complete, Q-type Cys(2)His(2)-zinc finger DNA-binding domain and a putative C-terminal ERF-associated amphiphilic repression (EAR) domain (Supplementary Figure 3). Zinc-coordinating histidine residues within the zinc finger domain are spaced by three amino acid residues. The C2H2 zinc finger and EAR motifs in PsPALM1a and PsPALM1b show no divergence. These motifs are further conserved in orthologous proteins from other legumes or even distant plant species including *Populus trichocarpa*, *Vitis vinifera*, and *Arabidopsis thaliana* (Supplementary Figure 3). Based on sequence data, *PsPALM1a* and *PsPALM1b* are likely to encode proteins with similar structures and functions even though the replacement of a conserved serine amino acid by a phenylalanine residue in PsPALM1a in an immediately adjacent site to the Zinc Finger domain is worth considering for any potential impact on the binding of the protein to DNA.

The 5’ upstream sequences of *PsPALM1a* and *PsPALM1b* were further examined and compared to the 5’ upstream sequence of *M. truncatula PALM1* (Supplementary Figure 4A), the only orthologous gene with a known function so far. The identity was low between *PsPALM1a* and *PsPALM1b* upstream regions when considering 2.5-kb sequences but reached 93 % in the first 495 bp upstream of the start codons. Three deletions of 8, 14 and 115 bp were observed in this interval in *PsPALM1b* compared to *PsPALM1a* (Supplementary Figure 4B). Like *M. truncatula PALM1* (Peng et al., 2017), several groups of putative cis-elements similar to the canonical auxin response element (AuxRE) TGTCTC are present in the promoters of *PsPALM1a* and *PsPALM1b*. In total, 19 and 13 TGTCXX elements were identified within the 2.5-kb 5’ upstream sequences of *PsPALM1a* and *PsPALM1b*, respectively. Some of these elements are conserved with *M. truncatula PALM1* or are shared exclusively between the 5’ upstream sequences of *PsPALM1a* and *PsPALM1b* (Supplementary Figure 4A; Supplementary Table 5).

**Supplementary Table 5.** See separate file. Positions of putative auxin response elements (AuxRE) in the 2.5-kb 5’ upstream sequences of *PsPALM1a*, *PsPALM1b* and *Medicago truncatula PALM1*. The canonical AuxRE, TGTCTC, is in bold. The positions where AuxRE cis-elements are conserved across the three sequences are indicated in bold green. The positions where AuxRE cis-elements are conserved across only two sequences are in bold orange. For details, please refer to Supplementary Figure 4.

**Figure S3.**
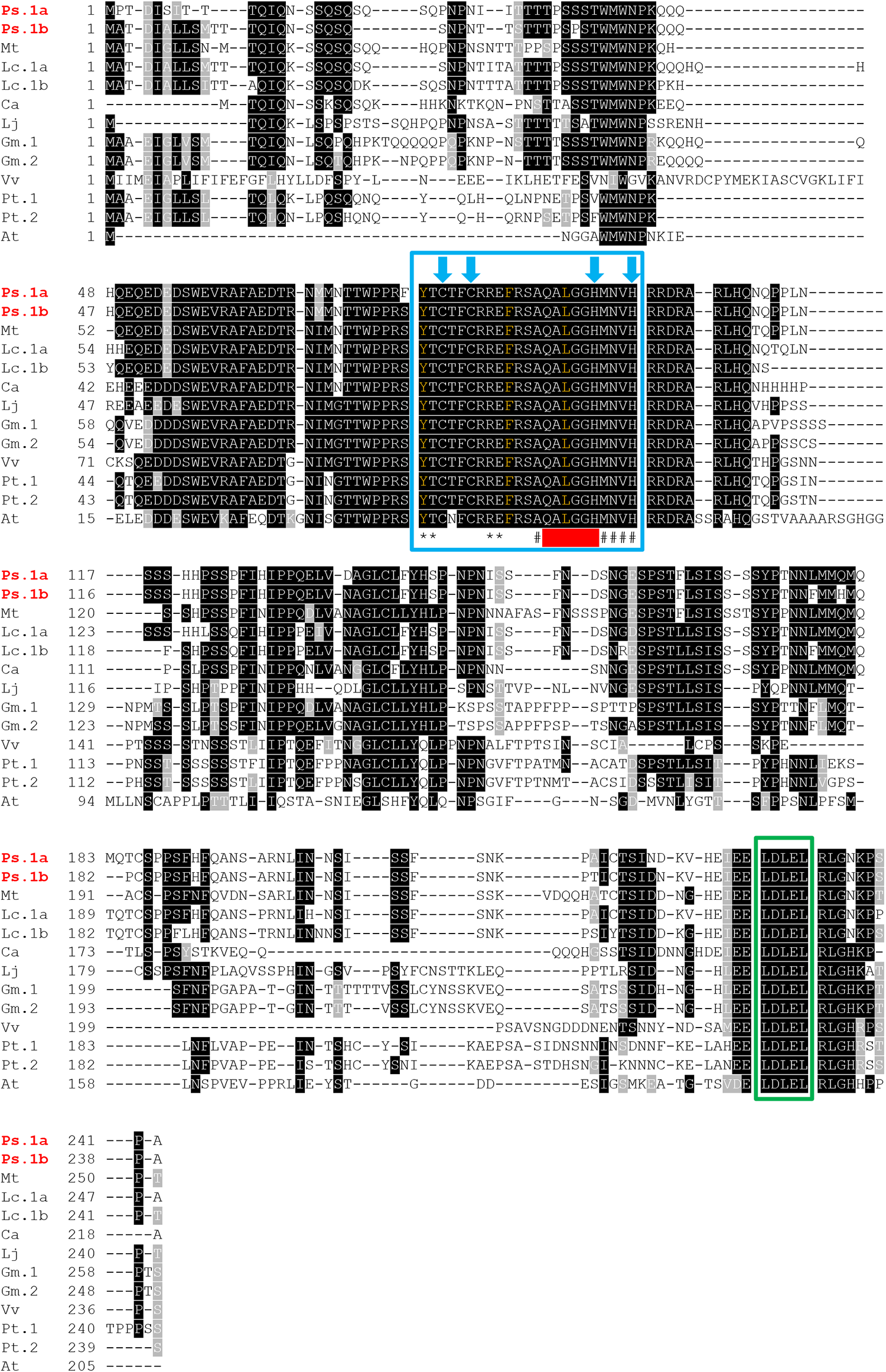
Alignment of predicted amino acid sequences of legume and non-legume genes orthologous to *Medicago truncatula PALMATE-LIKE PENTAFOLIATA1*. All sequences have a single Cys(2)His(2) zinc finger domain which is boxed in blue. Vertical blue arrows indicate the positions of conserved Cysteine and Histidine residues. Three amino acid residues separate the two Histidines. The invariant QALGGH motif, located inside the zinc-finger domain and characteristic of plant Q-type C2H2 zinc-finger proteins, is underlined in red. Hashes mark the amino acid residues that would compose, with the QALGGH sequence, an α-helix in the three-dimensional structure of the protein. Asterisks mark the residues that likely compose short β-sheets. Conserved Tyrosine, Phenylalanine and Leucine residues, which are expected to contribute, with the zinc ion, to hold together the α-helix and β-sheets (Wei et al., 2016) have their symbols written in orange. The core sequence of the Ethylene-responsive element binding factor-associated amphiphilic repression motif is highlighted by a green box. The correspondence between sequence IDs and gene IDs is as follows: - Ps.1a, *PsPALM1a*, *Psat2g173880*, chr2LG1:412902216-412902944; - Ps.1b, *PsPALM1b*, *Psat2g173360*, chr2LG1:412380009-412380728; - Mt, *PALM1/IRG1*, *MtrunA17Chr5g0400571*, MtrunA17Chr5:4836185-4836940; - Lc.1a, *Lc21900*, LcChr5:11776356-11777102; - Lc.1b, *Lc21107*, LcChr5:11841262-11841990; - Ca, *Ca9592*, Chr08:2811966-2812622; - Lj, *LjPALM*, *Lj2g3v1984740*, Chr02:29193053-29193808; - Gm.1, *GmPALM1*, *GLYMA_11G043200*, Chr11:3175603-3176385; - Gm.2, *GmPALM2*, *GLYMA_01G198700*, Chr01:53251395-53252147; - Vv, *GSVIVT01013168001*, chr02:7241430-7242586; - Pt.1, *PtrZFP56*, *Potri.001G142900*, Chr01:11626831-11627568; - Pt.2, *PtrZFP102*, *Potri.003G091200*, Chr03:11820160-11820879; - At, *AT4G17810*, Chr04:9906918-9907532. Ps *Pisum sativum*, Mt *Medicago truncatula*, Lc *Lens culinaris*, Ca *Cicer arietinum*, Lj *Lotus japonicus*, Gm *Glycine max*, Vv *Vitis vinifera*, Pt *Populus trichocarpa*, At *Arabidopsis thaliana*

**Figure S4.** See separate file. Distribution of auxin-response elements (AuxREs) in the 5’ upstream regions of *PsPALM1a*, *PsPALM1b* and *Medicago truncatula PALM1*. (A) Alignment of 5’ upstream, coding and 3’ downstream regions of *PsPALM1a*, *PsPALM1b* and *M. truncatula PALM1*. For all 3 genes, 2,500 bp and 500 bp long sequences were examined at the 5’ upstream and 3’ downstream regions, respectively. The ATG and stop codons are boxed in green. Sequences corresponding to putative AuxREs are in bold and colored. Red color is used for the AuxREs from *M. truncatula PALM1* that were already reported and described by Peng et al. (2017). Putative AuxREs for *PsPALM1a* and *PsPALM1b* are in blue. AuxREs that were reported to interact with *M. truncatula* AUXIN RESPONSE FACTOR3 (MtARF3) in tobacco leaves and significantly repress reporter luciferase gene expression (Peng et al. 2017) are underlined. AuxREs that did not exhibit significant inhibitory effects on luciferase activities in tobacco but were shown to be sufficient to be recognized by MtARF3 *in vivo* using chromatin immunoprecipitation coupled with polymerase chain reaction (Peng et al. 2017) are dotted underlined. The others were either not described in *M. truncatula* (TGTCAT, TGTCGG, TGTCGT) or have shown no interaction with MtARF3 (TGTCAC, TGTCCT). (B) Alignment of conserved promoter regions of *PsPALM1a* and *PsPALM1b*. The total length of the region is of 495 bp, based on data from *PsPALM1a*. The positions of AuxREs and the ATG stop codon are indicated.

### A fast-neutron mutant with a deletion of *PsPalm1a* but not *PsPalm1b* exhibits a palmate leaf phenotype in the JI2822 background

A line isolated from a fast-neutron-irradiated mutant population derived from accession JI2822 (Domoney et al. 2013) was identified by a forward genetic screen to exhibit an altered leaf phenotype. This line, named FN2070, exhibits extra leaflets at lower nodes and normal leaves at upper nodes (Supplementary Figure 5). Since pea Cameor v1a genome sequence (Kreplak et al. 2019) placed *PsPALM1a* and *PsPALM1b* 521 kb apart (Figure 5) most likely because of a misorientation of respective scaffolds, comparison of FN2070 genomic reads with the recently published genomic sequence of the leafy pea cv. ZW6 (Yang et al. 2022) having a higher assembly resolution at the *af* locus was undertaken. It showed a deletion encompassing multiple genes. The deletion concerns the consecutive genes receiving IDs *Psat2g173880* (*PsPALM1a*), *Psat2g173920* (PsCam002388; Figure 4) and *Psat2g173960* in Cameor’s v1a genome annotation, but not *Psat2g173360* (*PsPALM1b*). This confirms that the *PsPALM1a* genomic region is involved in leaf development and shows that the deletion of *PsPALM1a* (Chr2: 474040945-474041944) alone does not induce the af phenotype.

**Figure S5.**
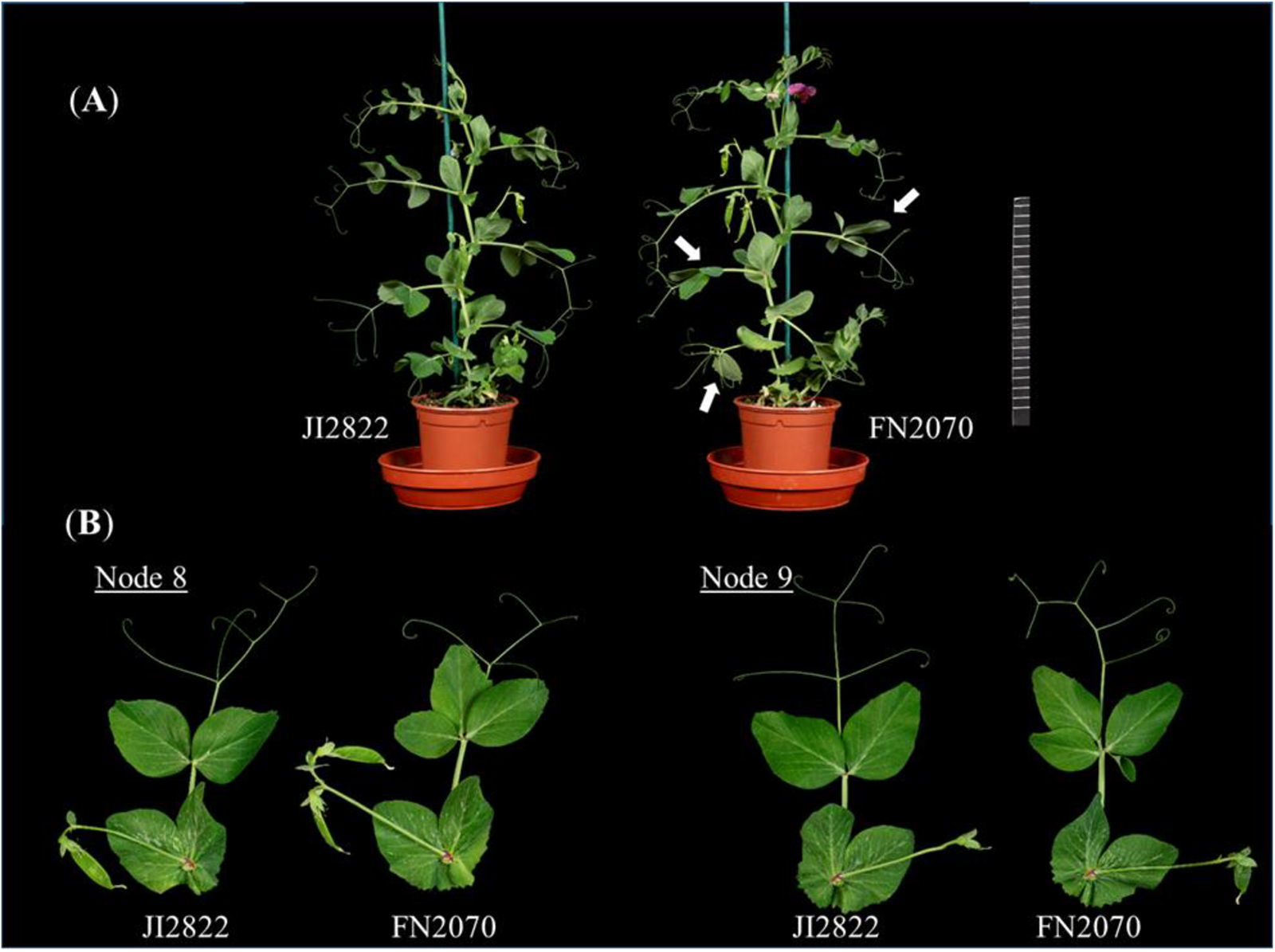
Fast Neutron line FN2070 and progenitor wild type accession JI2822. (A) Photos of 7-week-old plants of the fast neutron mutant (right) and the wild type accession (left). (B) Photos of detached leaves from the same plants as in A. Scale is in cm.

### TiLLING mutants of *PsPALM1a* or *PsPALM1b* in cultivar Cameor do not have the afila phenotype and no double mutant could be obtained

To further investigate whether *PsPALM1a* and/or *PsPALM1b* are involved in the regulation of compound leaf morphogenesis in pea, the Cameor ethylmethane sulfonate (EMS)-mutagenized population (Dalmais et al. 2008) was screened for mutants. No nucleotide substitutions introducing premature stop codons were found for *PsPALM1a*, however, the TILLING screen identified 15 missense mutations including two nonconservative amino acid changes in the C2H2 zinc-finger domain of the protein (Supplementary Table 6). Line 3715 has a change of Cysteine into Tyrosine at position 81 (C81Y) corresponding to the first conserved Cysteine residue of the zinc finger. Line 2512 has a H101Y substitution corresponding to the second conserved Histidine residue which occurs in a region that is expected to form an α-helix in the three-dimensional structure of this type of zinc finger (Wei et al. 2016). Both changes are expected to disrupt Zn^++^ ion binding within the zinc finger of the PsPALM1a protein and affect its DNA-binding function. In contrast, 13 missense and two nonsense mutations were identified for *PsPALM1b* (Supplementary Table 6). Translation termination would occur immediately before the QALGGH motif (Q91*) in line 2835 leading to a short protein with a partial DNA-binding C2H2 zinc-finger domain and no repressor domain (EAR motif). Unlike the palmate leaf phenotype exhibited by the *PALM1a* deletion in mutant FN2070 and the bipinnate *afila* leaf phenotype exhibited by the Bulgarian gamma-ray mutants lacking *PsPALM1b* (haplotypes 6 and 7, Figure 1E and Figure 4), all three EMS mutants in Cameor exhibited a conventional leaf phenotype (Supplementary Figure 6). F_1_ plants resulting from crosses between TILLING lines 3715, 2512 or 2835 and JI1195, a semi-leafless accession lacking both *PsPALM1a* and *PsPALM1b*, showed palmate or bipinnate leaf phenotypes at occasional nodes (Supplementary Figure 6), but most leaves were wild type in appearance. Bipinnate tendrilled leaves without leaflets, typical of *af* cvs. (Figure 1D), were not observed on any F_1_ plants. This showed that homozygous or hemizygous missense or nonsense alleles at either *PsPALM1a* or *PsPALM1b* did not confer an af phenotype. However, in hemizygous F_1_ plants some nodes produced bipinnate or palmate leaves.

**Figure S6.**
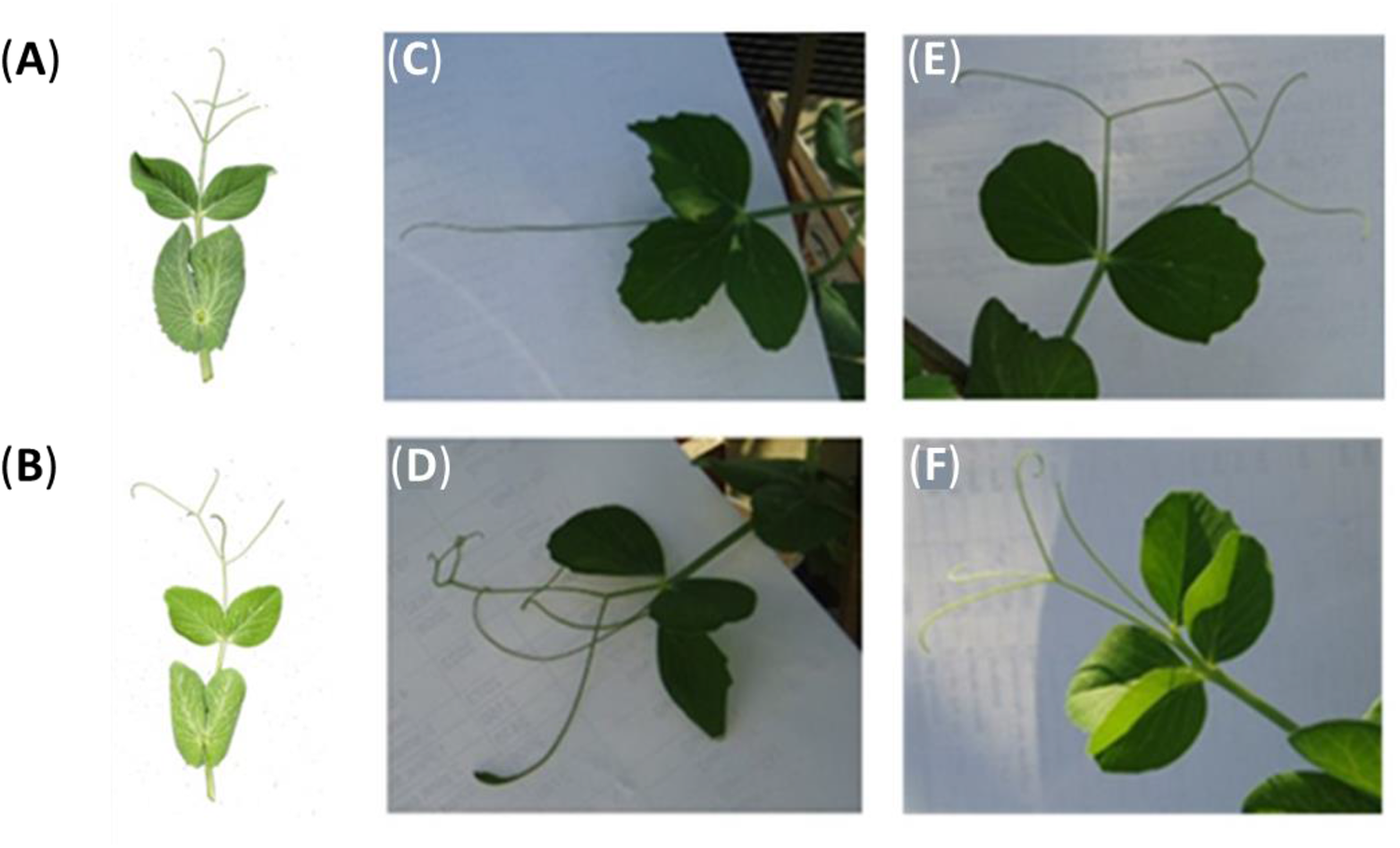
Photographs of adult pea leaves. (A) Conventional leaf from cultivar Cameor. (B) Leaf with wild type phenotype from EMS mutant in cv. Cameor, line 2385 (Q91*). (C) Leaf from an F_1_ line resulting from a cross between semi-leafless accession JI1195 and EMS mutant line 2512 (H101Y).

Double mutants could not be obtained by crossing *PsPALM1a* and *PsPALM1b* EMS TILLING mutants, because the recombination fraction between these is less than 0.43 cM. To further explore the proximity of these two genes, the Psa-B-Cam BAC library was screened (https://cnrgv.toulouse.inrae.fr/library/genomic_resource/Psa-B-Cam) with specific probes for *PsPALM1a* and *PsPALM1b*. Ten positive BAC clones were identified and validated by PCR (Supplementary Table 7) but none of these was positive for both genes. Electrophoresis data showed that the insert size of these BAC clones is relatively large: 74 (823E3), 80 (40C1) and 106 (490K14) kb and Sanger sequencing (Supplementary Table 7) suggested high abundance of transposon-related elements and repetitive sequences. This is congruent with pea genomic sequences becoming available later during the study. The genomic sequence of ZW6 (Yang et al. 2022), confirmed to a large extent by a recent investigation with Oxford Nanopore PromethION sequencing-based assembly of cv. Cameor (unpublished results), showed that *PsPALM1a* and *PsPALM1b* are separated by 259.6 kb. The lentil cv. CDC Redberry v1.2 genome assembly likewise showed two co-orthologs of *M. truncatula PALM1* (Figure 5, Supplementary Figure 7) but these are located about 64 kb apart with a single open reading frame located between them most likely related to a repeat element.

**Figure 5.**
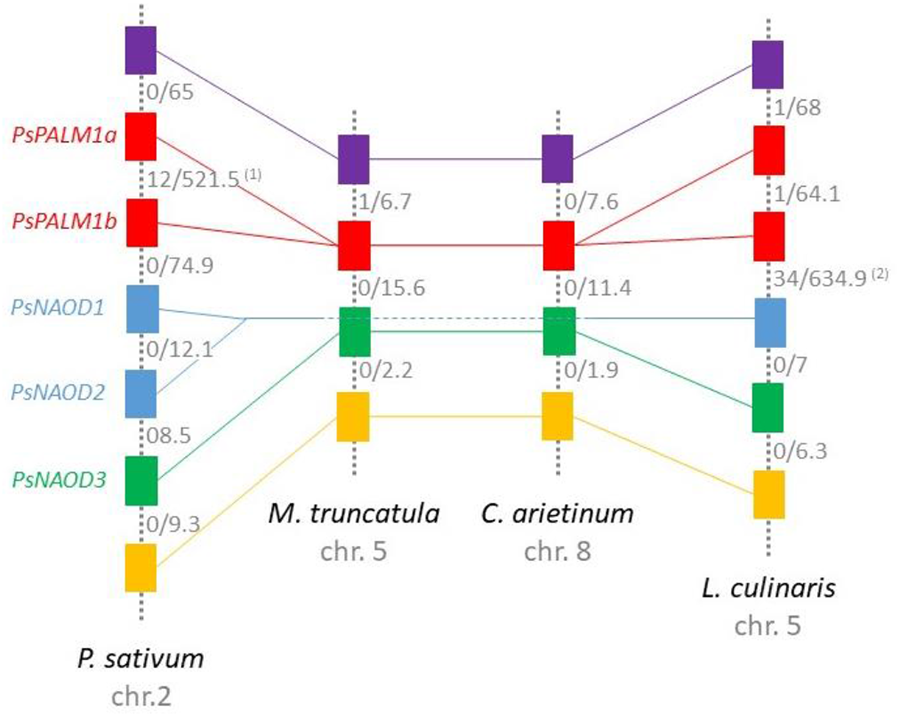
Evolution of the *af* containing region in the Fabeae tribe. Two representatives of the Fabeae tribe (*Pisum sativum* and *Lens culinaris*) and one representative of each of the sister tribes Trifolieae (*Medicago truncatula*) and Cicereae (*Cicer arietinum*) are included. Each rectangle refers to a different gene entity. Colors are used to distinguish the ortholog sets: in purple, putative small auxin-up RNA (SAUR) coding genes; in red, putative Q-type C2H2 zinc finger coding gene; in blue and green, putative N2-acetylornithine deacetylase coding gene; in orange, putative CAAX amino-terminal protease coding genes. Phylogenetic analyses were run to confirm orthology and the phylogenetic trees are available in fig. S7. Numbers in grey refer to the number of genes separating two illustrated gene entities (left) and to the physical distance in between (right; in Kb). The following genome assembly versions were used to construct this figure: *Pisum sativum* v1a, *Medicago truncatula* r5.0, *Cicer arietinum* Desi uwa-V3.0 and *Lens culinaris* CDC Redberry Genome v1.2.

**Figure S7.**
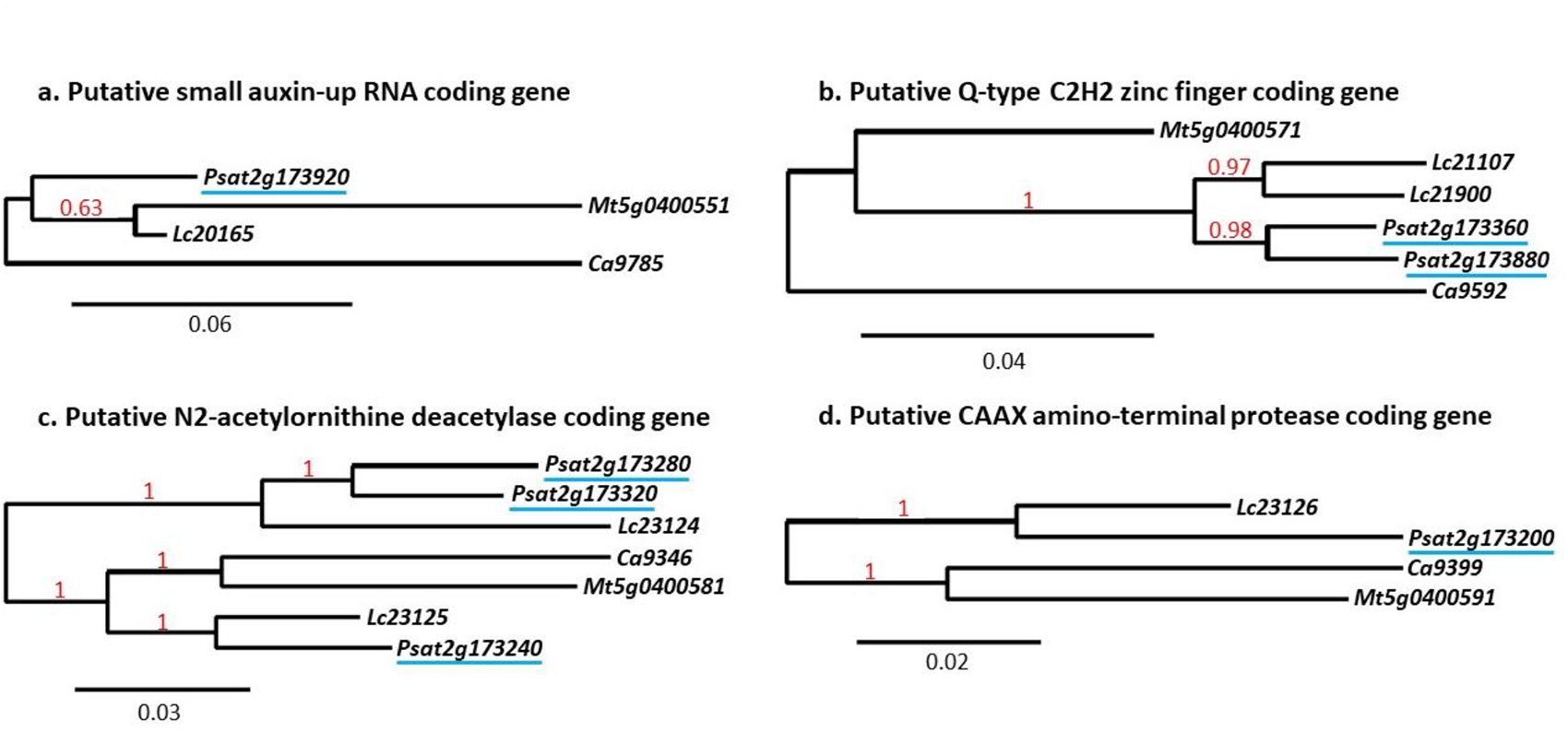
Phylogenetic trees constructed using the nucleotide sequences of the genes at the *af* locus. This supplementary figure was constructed to provide support for Figure 5 and highlight the orthology/homology of illustrated genes. Sequences are from: *Pisum sativum* v1a, *Lens culinaris* CDC Redberry v1.2, *Medicago truncatula* r5.0 and *Cicer arietinum* Desi uwa-V3.0 genome assemblies. The identifiers of pea genes are underlined. Phylogenetic trees were constructed using the Bayesian Inference method with the generalised time reversible (GTR) substitution model in Phylogeny.fr (Dereeper et al., 2008). Multiple sequence alignments were carried out using T-coffee (Notredame et al., 2000). Conserved blocks from multiple alignments were selected by using Gblocks (Castresana, 2000). Trees were implemented in MrBayes 3.2.6 (Huelsenbeck and Ronquist, 2001). Analyses were run for 100.000 generations starting from random trees, with trees sampled every 100 generations. The first 1000 trees were discarded as “burn-in”. The Bayesian inference trees were congruent with trees inferred with Neighbour-joining and Maximum likelihood methods (data not shown). (D-E) Leaves from F_1_ individuals resulting from a cross between EMS mutant line 3715 (C81Y) and JI1195. (F) F_1_ derived from a cross between mutant line 2385 and JI1195.

**Supplementary Table 6.** See separate file. EMS mutants for *PsPALM1a* and *PsPALM1b* in cv. Cameor identified using a TiLLING approach. Only lines with missense or nonsense mutations are included in this table. The lines that are predicted to have the most severe disruption to the encoded proteins are highlighted in green.

**Supplementary Table 7.** See separate file. Positive BAC clones for PsPALM1a and PsPALM1b and analysis of BAC-end sequences (BES).

### Virus-induced gene silencing did not confer the afila phenotype

Since the physical proximity of *PsPALM1a* and *PsPALM1b* hinders the construction of a double mutant, virus-induced gene silencing (VIGS) was attempted as a reverse genetics tool to test the functions of *PsPALM1a* and *PsPALM1b*. VIGS is a suitable strategy for knocking down both *PsPALM1a* and *PsPALM1b* because of their high degree of sequence similarity and this was attempted with a Bean pod mottle virus (BPMV) VIGS vector (Zhang et al. 2010; Meziadi et al. 2016). We used cv. Champagne (Supplementary Figure 8), a forage pea with conventional leaf type. The BPMV-*PsPALM*1 VIGS vector was delivered to pea plants at the two-leaf stage of development by mechanical rubbing of stipules and leaflets of both leaves. At maturity, no leaves with a clear *afila* phenotype were observed.

Pea leaves are heteroblastic whereby additional pinna pairs are added as the plant matures. This increase in complexity is seen first as an additional pair of tendrils and then as an additional pair of leaflets (Blixt 1972). Occasionally, a leaflet forms opposite a tendril as this increase in pinna number occurs. We noticed a slight increase in these two leaf forms (an additional tendril pair or a leaflet opposite a tendril) in the VIGS treated vs control plants (35 out of 352 leaves in the BPMV-*PsPALM1* compared to 2 out of 85 in the BPMV-0-inoculated plants; χ^2^= 5.09; p = 0.024) (Supplementary Figure 8). These results suggested that *PsPALM1a* and *PsPALM1b* are involved in the control of leaf formation and that a full knockout of both genes is high likely needed to obtain an af phenotype.

**Figure S8.**
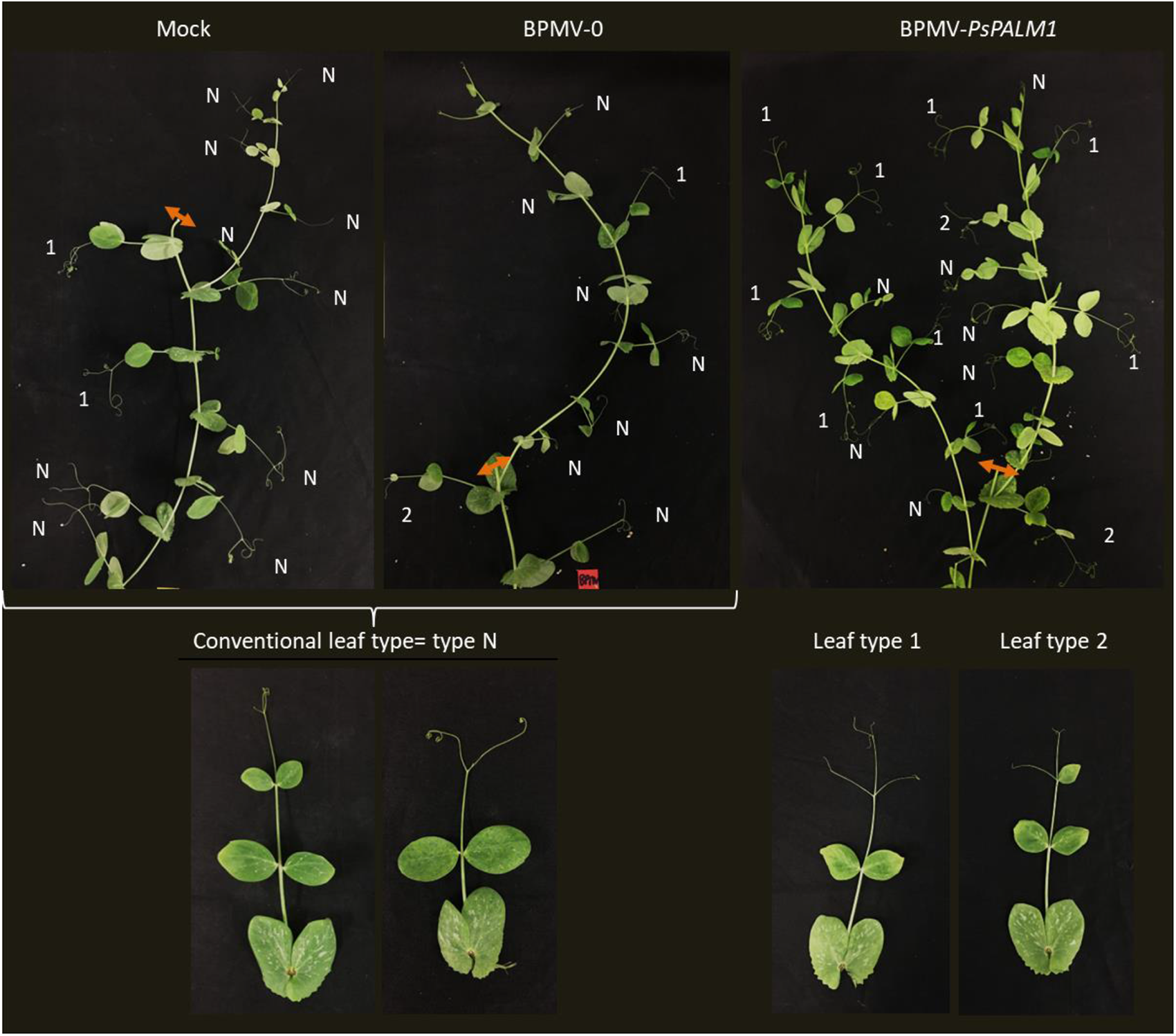
Attempted *PsPALM1* virus-induced gene silencing in conventional leafy *Pisum sativum* cv. Champagne. Bean pod mottle virus BPMV was used as a vector to knock-down the expression of *PsPALM1a* and *PsPALM1b*. Representative stems of plants inoculated with buffer (Mock, left panel), BPMV-0 (middle panel), and BPMV-*PsPALM1* (right panel) were photographed under natural light at 11.5 weeks post-inoculation (wpi). Pruning of the stems was done at 4 wpi and the site of cutting is shown with an orange arrow. Leaf morphologies were scored and classified in 3 types: type N (1 pair of stipules +1-2 pair(s) of leaflets +2-3 tendrils), type 1 (1 pair of stipules +1 pair of leaflets + 2 or more pairs of tendrils and a terminal tendril) and type 2 (with a leaflet opposite to a tendril). The combined occurrence of type 1 and 2 leaves is significantly higher in BPMV-*PsPALM1* silenced plants.

### *PsPALM1a* and *PsPALM1b* exhibit similar expression profiles and contribute to the regulation of leaf morphology in a complex, likely dose-dependent, manner

Microfluidic droplet-based gene expression profiling was conducted to study the expression of *PsPALM1a* and *PsPALM1b* at the spatio-temporal level. RNA extractions were performed from root, nodule, shoot, leaf and seed tissues of cv. Cameor (Supplementary Figure 9). No expression was detected for *PsPALM1a* or *PsPALM1b* in shoot, root or nodule samples. Expression of both genes was detected mainly in developing seeds 12 days after pollination (DAP) and in germinating seeds. Expression was detected in the embryo and not in the albumen or the seed coat at this particular stage of seed development (Supplementary Figure 9) when leaf primordia are developing on the embryonic axis. These results are in line with the data from the pea gene atlas (Alves-Carvalho et al. 2015) visible on the Pea genome browser (https://urgi.versailles.inra.fr/Species/Pisum) for genes *Psat2g173880* and *Psat2g173360* (Kreplak et al. 2019) corresponding to *PsPALM1a* and *PsPALM1b*, respectively. The expression of *Uni* was lower than that of *PsPALM1a* and *PsPALM1b* in all tissues except for the embryo (Supplementary Figure 9).

**Figure S9.**
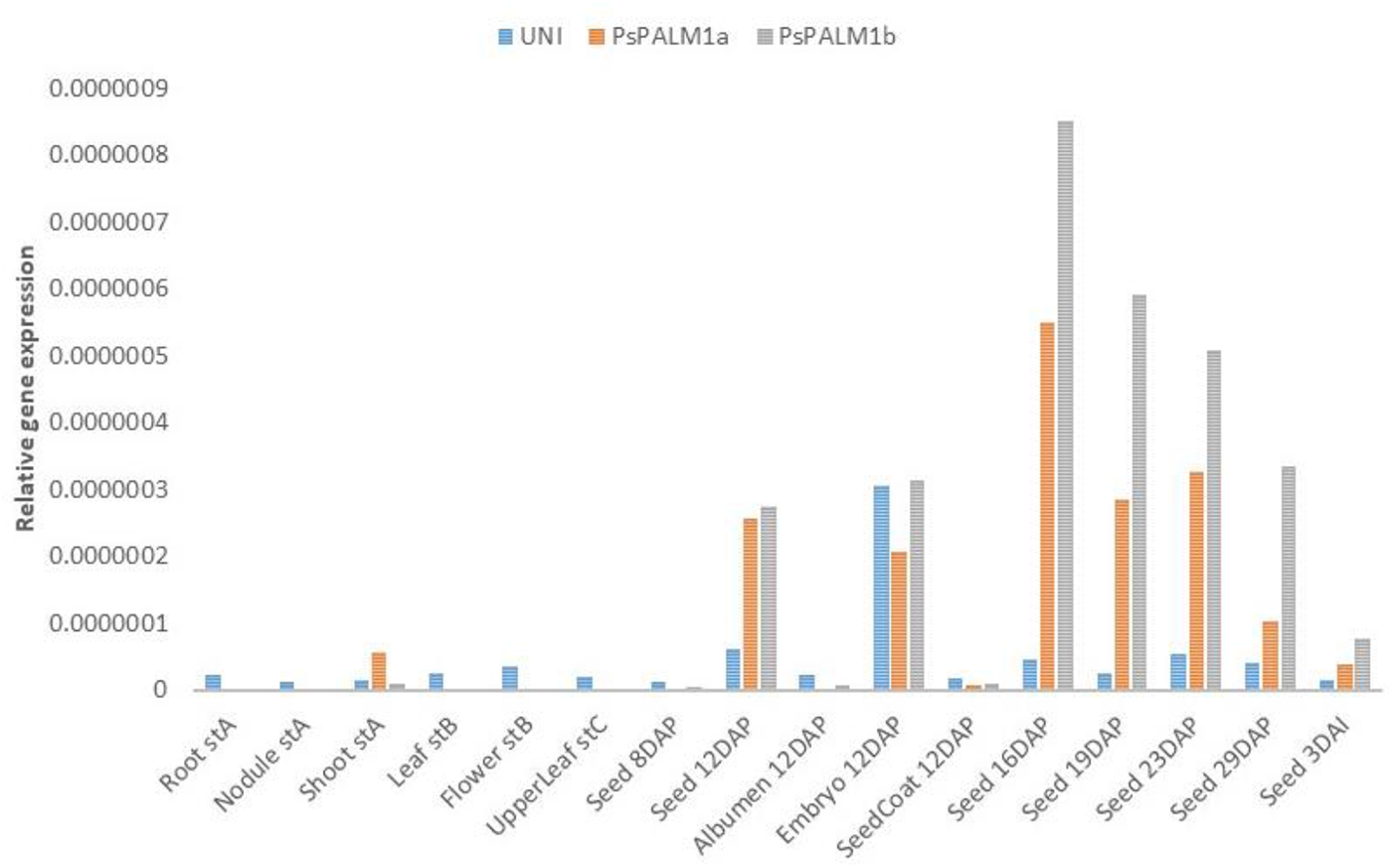
Expression of *UNI, PsPALM1a* and *PsPALM1b* in different tissues under normal growth conditions detected by qRT-PCR. stA=7–8 nodes, 5–6 opened leaves; stB=beginning of flowering stage; stC=20 days after the beginning of flowering. DAP=days after pollination; DAI=days after imbibition.

The expression of *PsPALM1a* and/or *PsPALM1b* was further studied in the context where *PsPALM1b* harbours a mutation provoking a premature stop codon (EMS TILLING mutant 2835 in the Cameor background) or is deleted (Bulgarian gamma-ray mutants in cvs. Auralia, Borek, and Raman). Ten germinating seeds at three days post-imbibition were considered per condition with each being a separate biological replicate. The expression level of *PsPALM1a* and *PsPALM1b* did not significantly change in the EMS mutant compared to the Cameor wild type (Figure 6A). However, the expression of *PsPALM1a* in the Bulgarian gamma-ray mutants significantly decreased (Figure 6B-D). The mutant in cv. Raman (also known as 11/47, Ambrose 2004) exhibited the least decrease (Figure 6D). This differential regulation of *PsPALM1a* might explain the divergent phenotypes observed in these mutants, with the TILLING mutant 2835 having a normal leaf architecture, the mutation in Bulgarian line 11/47 (JI 3129) provoking an intermediate phenotype (Figure 1E) and the mutations in Bulgarian lines 1/534 and 2/544 resulting in bipinnate tendrilled leaves (Supplementary Figure 2).

**Figure 6.**
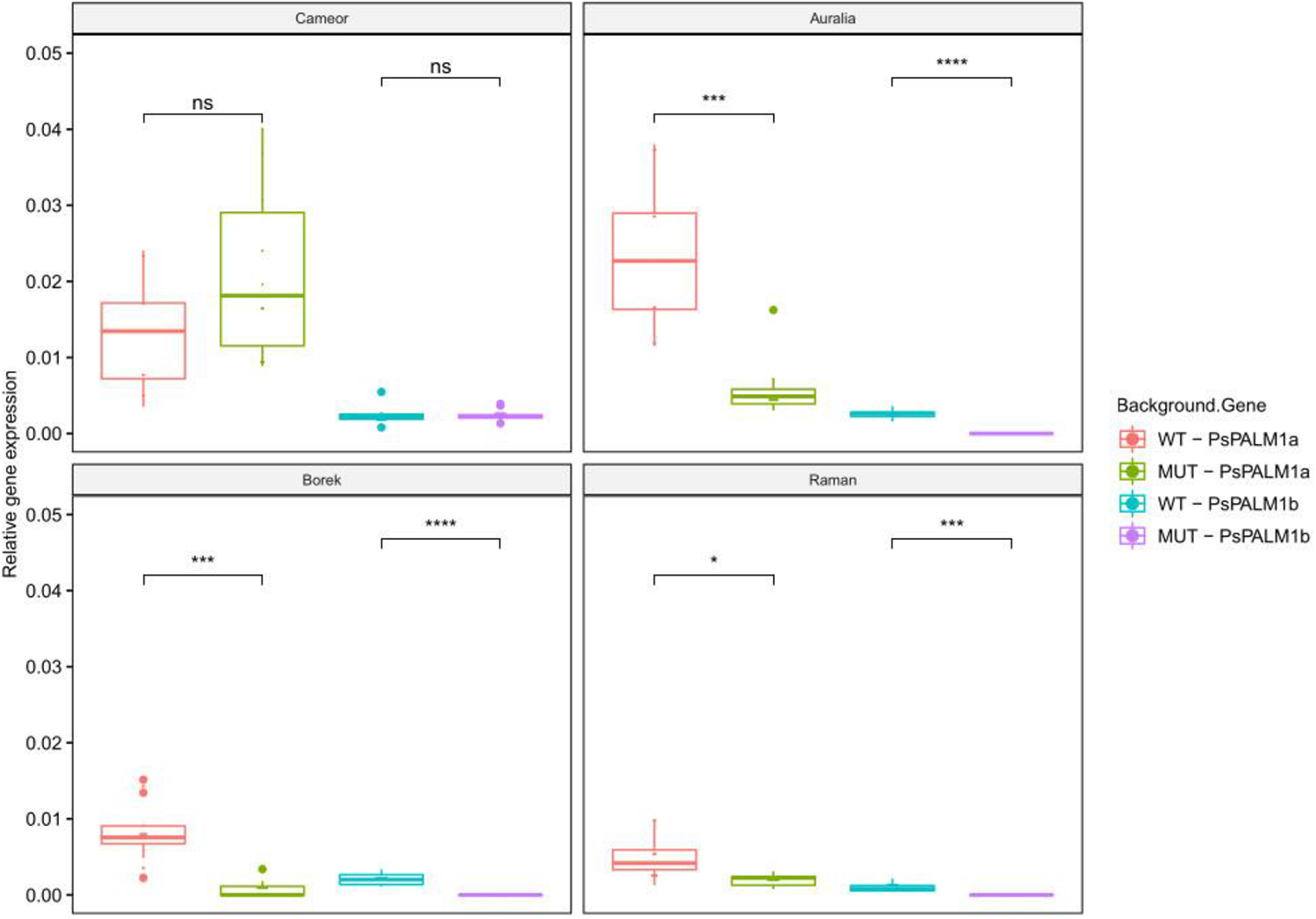
Expression of *PsPALM1a* and *PsPALM1b* in germinating seeds of four pea cultivars and their respective mutants. Gene expression was detected by qRT-PCR. The pea cultivars are Cameor, Auralia, Borek and Raman. All mutants carry mutations at *PsPALM1b*. The mutant in Cameor background was identified in an ethylmethane sulfonate (EMS)-mutagenized population (line 2835) and has a G:C to A:T transition leading to a premature stop codon. The mutants in the other backgrounds have full deletions of *PsPALM1b* that arose from mutagenesis with gamma radiation. Seeds were sampled at three days post-imbibition and ten biological replicates were used per condition. Graphics and statistics were done with the software R V.4.0.0 using the ggpubr package V.0.3.0, available at https://github.com/kassambara/ggpubr. ns, p > 0.05; *, p <= 0.05; **, p <= 0.01; ***, p <= 0.001; ****, p <= 0.0001, where p = P-value using paired t-test.

## Discussion

Fifty years ago, the pea crop was different from today. Plants were tall and had conventional leaves, constituted of stipules, leaflets, and tendrils. This often resulted in a heavy canopy with poor standing ability. Because of lodging, much of the yield was lost at harvest and the quality of the seeds was often poor due to pathogen attacks arising from damp soil conditions (Banniza et al. 2005). The success story of improving standing ability in pea owes a great deal to the introduction of the *af* mutation (Kujala 1953; Solov’eva 1958; Goldenberg 1965; Khangildin 1966; Swiecicki 1982) together with a mutation in Mendel’s stem length gene *Le* (White 1917; Lester et al. 1997; Martin et al. 1997) into breeding material. The *af* mutation results in leaves with an increased number of tendrils. Each af leaf produces lateral, transiently indeterminate, primordia in place of leaflet primordia. These primordia become pinnae, which in the presence of the dominant *Tl* allele, produce lateral tendrils and terminate as tendrils. The *le* mutation causes short stem internodes (Martin et al. 1997; Lester 1997). The major scientific and breeding efforts towards achieving a short-stem semi-leafless ideotype were undertaken in the seventies especially in Europe (Duparque 1996; Mikić et al. 2011b). Today’s dry pea cvs. almost all carry the *af* mutation, and the recessive *le* allele.

Here we identify the *af* mutations as large deletions encompassing at least 450 kb on pea chromosome 2. Such structural variants are known to contribute to genetic diversity and phenotypic variation in living organisms (Hurles et al. 2008; Springer et al. 2009) and these are often associated with transposable elements, DNA recombination, replication, and repair-associated processes (Morgante et al. 2005; Hastings et al. 2009). In rice, a 38.3-kb fragment harbouring genes regulating grain number, plant height and the gene *heading date7*, is completely deleted in the cv. Zhenshan 97 (Xue et al. 2008). A 254-kb genomic deletion reduces palmitic acid level in soybean seeds (Goettel et al. 2016). In maize, the deletion of a 5.16-Mb segment on chromosome 2 in the spontaneous *multi-trait weakened* mutant leads to a loss of 48 genes which is responsible for a set of phenotypic defects, including wilting, decreased yield, reduced plant height, increased stomatal density, and accelerated water loss (Han et al. 2019). Pea has a genome of ca. 4.45 Gb, an estimated 83% of which comprises repetitive elements, mostly retrotransposons (Kreplak et al. 2019). The presence or absence of these elements across the genome is known to be highly polymorphic among pea germplasm collections. The presence/absence polymorphism associated with retrotransposons may sometimes be accompanied by presence/absence of the sequence between adjacent retrotransposons and so contribute to genic diversity (Ellis et al. 1998; Jing et al. 2010).

Seven haplotypes were identified at the *af* locus in the pea accessions exhibiting an af phenotype in this study, namely haplotypes 2 to 8 (Figure 4 and Supplementary Table 4). The assignment of 107 semi-leafless (*afaf StSt*) and leafless (*afaf stst*) pea accessions corresponding to cvs., mutants and research and breeding lines into these seven haplotype groups is broadly congruent with available pedigree information, reflecting the robustness of these molecular tools to discriminate the different haplotypes.

The cv. Ballet, selected from a cross between Filby x NRPB 60242 (Leterme et al. 1992), and its derived RILs (Pop4-69, Pop5-22, Pop5-80, Pop5-81, Pop5-142; Supplementary Table 4) all clustered, as expected, with Filby. The varieties Sum (Porta x Wasata), Hamil ([(Wasata x 1.6L/78) x Porta]) and Ramir ([(Porta x Wasata) x Flavanda]) all have in their pedigree the induced mutant in cv. Wasata (Jaranowski 1970) obtained in Poland (FAO/IAEA Mutant database; https://mvd.iaea.org/) and are clustered accordingly.

Inconsistencies can be seen (Supplementary Table 4), but most likely these highlight inaccuracies in the information sources, or records because until now, *af* alleles have been indistinguishable. For instance, contrary to the records of the FAO/IAEA Mutant database indicating that the *af* allele in Legenda derives from spontaneous mutation of cv. Usatyj, Legenda and Usatyj-*af* were found in two different haplotype clusters (Supplementary Table 4), but this is in agreement with Kielpinski and Blixt (1982) and reported by Kielpinski in 2017 (http://www.kielpinski.eu/). Legenda is similar to cvs. Sum, Hamil, Ramir, and Miko, all developed through hybridization with Wasata. Also, the records of the John Innes *Pisum* collection (http://bioinf.scri.ac.uk/germinate_pea/app/) indicate that the first leafless cv. in the United Kingdom (Ambrose 2004), registered for Plant Variety Rights in 1977, viz. cv. Filby (accession ID: JI 1768), derives from a cross between cv. Dik Trom and a spontaneous *af* mutant from Argentina. The Filby *af* allele clusters with the *af* used by Marx in lines B268-394-3, MISOG1 and MISOG2 and this is the Argentinian *af* mutant from Goldenberg (1965) according to handwritten breeding notes. However, the lines JI2400 and NGB101692, from a common origin, are recorded as carrying Goldenberg’s *af*, but in fact they carry the *af* allele from Solov’eva (1958): it therefore seems likely that these lines are mis-identified. Future complete sequencing of all pea accessions will shed more light on these inconsistencies.

The different haplotypes at the *af* locus show that several progenitors have been used for the development of improved pea cvs., since the first initiatives to breed semi-leafless and leafless pea cvs. launched in Poland and England (Snoad 1981; Kielpinski and Blixt 1982). Of the 77 *af* cvs. we examined (Figure 4; Supplementary Table 4), 6 derive from the spontaneous mutant reported by Solov’eva (1958) and 51 from Goldenberg (1965). The induced mutant in Wasata (Jaranowski 1970) is represented in 15 cvs. Five cvs. have *afila* alleles of unknown origin.

We propose that once the first cvs. became available, breeding programmes used these as the source material for constructing new breeding lines and these have been widely distributed throughout the world. Early examples were the use of the Russian and Argentinian alleles in the first *af* breeding programme in the UK. More recently, cooperative exchanges of plant material for breeding purposes occurred in Canada, where the progenitors of many semi-leafless Canadian varieties are lines from Svalöf Weibull, one of the largest plant breeding and seed groups in Europe, based in Sweden (Vandenberg et al. 2004; Warkentin et al. 2005).

Haplotypes 2, 7, and 8 (Figure 4) lack the three *PsNAOD1-3* genes. Molesini et al. (2015) have shown that *NAOD* contributes to the regulation of polyamine and ornithine levels in *A. thaliana* and that a reduced *NAOD* expression results in alterations at the reproductive level, causing early flowering and impaired fruit setting. It would be interesting to investigate if a poorer fitness of mutants carrying a deletion of *PsNAOD* genes in pea was similarly encountered and affected any attempts to use the mutants from haplotypes 7 and 8. Interestingly, an experiment using *af* near-isogenic lines of cv. ‘Terese’ (haplotype 3, lacking *PsNAOD1*) showed that carrying *af* was associated with a lower seed weight (Burstin et al. 2007).

All cvs. carrying the *af* mutation share a deletion that includes two genes on pea LGI or chromosome 2, viz. *PsPALM1a* and *PsPALM1b*. After *PALM1* in *M. truncatula* (Chen et al. 2010) and *St* in pea (Moreau et al. 2018), this represents further evidence for the involvement of Q-type C2H2 zinc finger transcription factors in the control of leaf morphology in legumes belonging to the inverted-repeat lacking clade. In both simple- and compound-leaf species, the *class I KNOTTED1-like homeobox* (*KNOXI*) genes play pivotal roles in the maintenance of meristem activity in the shoot apical meristem (Lincoln et al. 1994; Kerstetter et al. 1997; Lenhard et al. 2002). Downregulation of *KNOXI* genes occurs at the incipient sites of leaf primordia and the expression disappears in the meristem cells that are destined to be the next leaf (Smith et al. 1992; Lincoln et al. 1994; Long et al. 1996). In some compound leaf species, including *Solanum lycopersicum* and *Cardamine hirsuta*, the *KNOXI* genes are reactivated in developing leaf primordia and this reactivation is required for compound leaf development in these species (Bharathan et al. 2002). However, in *M. truncatula* and pea, *KNOXI* genes are repressed by *Crispa*, the pea orthologue of *Phan/AS1*, at the inception of the leaf primordium (Tattersall et al. 2005) and thus not shown to be involved in the early patterning of pea or *M. truncatula* compound leaves (Hofer et al. 2001; Champagne et al. 2007). *SGL1* in *M. truncatula* and *Uni* in pea, both orthologs of *A. thaliana LEAFY*, function in controlling leaf development and are required for the initiation of leaf lateral primordia (Hofer et al. 1997; Champagne et al. 2007; Wang et al. 2008).

*PALM1* negatively regulates the expression of *SGL1* in *M. truncatula* lateral leaflet primordia (He et al. 2020) and is in turn the direct target of *M. truncatula* AUXIN RESPONSE FACTOR3 (MtARF3) that acts as a transcriptional repressor of *PALM1* through interaction with specific AuxREs in the *PALM1* promoter (Peng et al. 2017). *M. truncatula* fast neutron mutants (*palm1-1* and *palm1-2*) having large deletions encompassing up to eight annotated genes have been described. Mutants carrying deletions (*palm1-3*) or tobacco Tnt1 retrotransposons (*palm1-4*, *palm1-5*, and *palm1-6*) within the coding region of *PALM1* also contributed to confirm its role as a determinacy factor for leaf morphogenesis in *M. truncatula* (Chen et al. 2010). In this study, we show that the absence of orthologs of *PALM1* confers the bipinnate tendrilled phenotype in pea, which indicates that the *PALM1* genes act to limit lateral morphogenesis in the pinnate wild type pea leaf.

*PALM1*, *PsPALM1a* and *PsPALM1b* have largely similar gene and protein structures and all harbour AuxRE cis-elements in their promoters. Similarity is also evident in their expression profiles. Like *PALM1* (Chen et al. 2010), *PsPALM1a* and *PsPALM1b* are expressed in developing seeds, but remain barely detectable in other tissues including roots, stems, flowers, pods, and seed coats. Detailed molecular investigations are needed to understand and compare the mode of function of these genes from pea and *M. truncatula*. The genetic models proposed for *Af* (Gourlay et al. 2000), the physiological studies by DeMason et al. (2013) and the investigations of *PALM1* by Peng et al. (2017) are predictive of conserved functions. Current models link auxin signalling to compound leaf patterning and highlight a pivotal role for *PALM1* or *Af* in this process. Now that *PsPALM1a* and *PsPALM1b* are identified and the ortholog of *MtARF3* in pea is known through the current genome assembly (*Psat1g202240*; Kreplak et al. 2019), the module *PsPALM1a*/*PsPALM1b*/*UNI*/ *Psat1g202240* can be dissected at the molecular level. The impact of *Af*, i.e. *PsPALM1a* and *PsPALM1b*, on the size of the stipules as suggested by Kielpinski and Blixt (1982) and any possible pleiotropic effects of these genes such as the role of *PALM1* as an inhibitor of rust germ tube differentiation (Uppalapati et al. 2012) also deserve to be studied.

To conclude, this study shows the causal link between the *af* mutation and a deletion of a genome region containing two *PALM1 g*enes in pea, and contributes to understanding the control of compound leaf morphogenesis in legumes. However, the mechanisms underlying the afila phenotype and the respective roles of both genes, the surrounding region and the genetic background need to be further examined. Nevertheless, it paves the way towards engineering new pea ideotypes where useful features could be brought by new *af* alleles or information-based management of breeding schemes. It is important to understand the evolutionary dynamics in the *af* region in the tribe Fabeae, which led to a tandem duplication of *PALM1* genes in pea and lentil. Translational knowledge can be applied now to engineer improved cvs. in related crops such as *L. culinaris* (lentil) and *V. villosa* (hairy vetch), for which lodging is a major production problem.

## Acknowledgments

The authors are grateful to Julie Fievet for her help with the statistical analyses of the VIGS experiments. They would like to acknowledge Karen Boucherot and Emilie Vieille for their contribution to molecular analyses (qPCR and genotyping), Sophie Blanchet for the construction of the BPMV-*PsPALM1* VIGS vector and Hélène Berges, Arnaud Bellec and Nathalie Rodde from the French Plant Genomic Resources Center for providing the BAC clones and evaluating their insert size. The authors would like also to thank the Germplasm Resources Unit at the John Innes Centre, and especially Mr. Mike Ambrose, the Nordic Genetic Resource Centre (NordGen) and the INRAE CRB PROTEA for providing seeds. A special thank you to the late Rosemary Harvey who translated Kujala’s 1953 publication into English. Marianne Chabert-Martinello, Céline Rond and Anthony Klein provided precious passport and phenotypic data related to accessions from INRAE CRB PROTEA and Sarah Wilmot assisted in retrieving archival records of the *afila* breeding programme at the John Innes Institute.

## Funding

French National Research Agency grant ANR-09-GENM-026, GENOPEA

French National Research Agency grant ANR-11-BTBR-0002, PeaMUST

UKRI Strategic Programme grant BBS/E/J/000PR799

European Union FP6 Framework Program FOOD-CT-2004-506223

John Innes Centre Strategic Fellowship (NE)

## Author contributions

Conceptualization: NT, NE

Methodology: NT, JH, GA, FJ, LT, PP, MD, CL, JK, SP, VG

Investigation: NT, JH, GA, NE

Funding acquisition: NE, JB

Supervision: NT, NE, JB

Writing – original draft: NT, NE

Writing – review & editing: NT, JH, GA, SP, VG, NE, JB

## Data S1

### Parallel approach to identify candidate genes for the af phenotype

Two independent studies in two independent laboratories were at first carried out to place *af* on pea genetic maps/genomic scaffolds and identify candidate genes. The approach relying on the four RIL populations, namely Pop2, Pop4, Pop5, and BxP, and on synteny between close legume genomes is described in the main text. The other approach that used JI813 x JI1201 RIL population and pea bacterial artificial chromosome (BAC) clones is described in this document. A recombinant inbred population of 58 RILs, derived from the cross between JI1201 and JI813 (from the John Innes Pisum germplasm), segregating for leaf type, was genotyped by cDNA-AFLP. In total, 332 segregating markers could be scored. Markers that mapped close to *af* were identified. A further 967 F_7_ and F_8_ individuals, segregating for *Af/af* were obtained by selfing an F_4_ heterozygote line. These individuals were genotyped using a combination of phenotypic markers (af and i) and linked genetic markers (GdcH and cDNA573; Supplementary Table 3). No recombinants were detected between GdcH and *af*. Within this region, two copies of a C2H2 Zn-Finger transcription factor encoding gene were found, which were present in *AfAf* genotypes and absent from homozygous *afaf* genotypes. These two genes could not be separated in the expanded 967 segregants obtained from a single af, heterozygote F_4_ line presumably because they both lay within a deletion in JI1201 (*af*) with respect to JI813 (*Af*).

Two BAC clones, 1041E7 and 1018I23, were identified from a BAC library of *Hin*dIII-digested cv. Cameor genomic DNA in pIndigoBAC5 that carried the *PsPALM1a* and *PsPALM1b* genes separately (Supplementary Table 7). The BAC library is hosted at the French Plant Genomic Resources Centre (CNRGV). Their coding sequences were found to be 93% identical. PCR primers specific to each gene were designed and used for segregation analysis in the *P.sativum* x *P.abyssinicum* RIL population JI2822xJI2022 (both are *AfAf*). Among these 135 F6 RILs, no recombinants were found between the two markers.

