## Supplementary Figure 4 for "*afila*, the origin and nature of a major innovation in the history of pea breeding"

**Figure S4.** Distribution of auxin-response elements (AuxREs) in the 5’ upstream regions of *PsPALM1a*, *PsPALM1b* and *Medicago truncatula PALM1*. (A) Alignment of 5’ upstream, coding and 3’ downstream regions of *PsPALM1a*, *PsPALM1b* and *M. truncatula PALM1*. For all 3 genes, 2,500 bp and 500 bp long sequences were examined at the 5’ upstream and 3’ downstream regions, respectively. The ATG and stop codons are boxed in green. Sequences corresponding to putative AuxREs are in bold and colored. Red color is used for the AuxREs from *M. truncatula* *PALM1* that were already reported and described by Peng et al. (2017). Putative AuxREs for *PsPALM1a* and *PsPALM1b* are in blue. AuxREs that were reported to interact with *M. truncatula* AUXIN RESPONSE FACTOR3 (MtARF3) in tobacco leaves and significantly repress reporter luciferase gene expression (Peng et al. 2017) are underlined. AuxREs that did not exhibit significant inhibitory effects on luciferase activities in tobacco but were shown to be sufficient to be recognized by MtARF3 *in vivo* using chromatin immunoprecipitation coupled with polymerase chain reaction (Peng et al. 2017) are dotted underlined. The others were either not described in *M. truncatula* (TGTCAT, TGTCGG, TGTCGT) or have shown no interaction with MtARF3 (TGTCAC, TGTCCT). (B) Alignment of conserved promoter regions of *PsPALM1a* and *PsPALM1b*. The total length of the region is of 495 bp, based on data from *PsPALM1a*. The positions of AuxREs and the ATG stop codon are indicated.

```
(A)

PsPALM1a    1 ACCATTTTTCGGAAGAGAACTC-T-----------GATTCTGATTTCTATGCTTAAG-GA
PsPALM1b    1 AGAAATA---------GA-TTT-TACC---TCGATTTTTAGTAATAACAAGGTAAAA-TA
MtPALM1     1 GATATCT---------AATTTTTTAATTATTTAATGAATAAAAACAACAAGCTTAACATG


PsPALM1a   48 ATCA---TCCATGTGTTGTTTCTTTC----GATTGCTTGTAACT---CTTCAACCATA-G
PsPALM1b   46 TTTT-------ATTGGAACATATTTTTTGAGATAAAATATAA-----AG---GTGTTTTT
MtPALM1    52 TTTAGCCCGCAAGTGAAGTATACTTA----ATTAACTAATACTAGGTAT---GAGTTA-G


PsPALM1a   97 CAGACTTCCACTTCATATTGTTTAAAGCTTC-ACTATAGTTGATTGG--TTCAACACCTG
PsPALM1b   91 TCAACGTC-----------CTTAAACAAATC-TTTGTCGTCCATTTGAGTATCACGCTTC
MtPALM1   104 CTAGCATC-----------CCTGTCTGTGAATTTGGTGGCATGTACTAGTTGGACGTCGA


PsPALM1a  154 C--AAGTAAAGCAAAA--TGAACTAATTATC-CACCTGGTATGACTTCATCATCACCAGT
PsPALM1b  139 A--AAACACTGCAAATAGTGATCT--TTGTGAAACCTA--ACGATCTCATTAT-AAAAGA
MtPALM1   153 CATAAACAATGATACA--GGAACG---TAAGAAAGGT---GTGTTTCTTTCTTTCTTGGT


PsPALM1a  209 CACTTCGTAGTCTTG-AAGTCTTGTTGGACGAACT--CCAGTTCTTTGAGGT--CTCTGA
PsPALM1b  192 -ATTTTGAATA-TTA-GTGAATCCCT-GA--AACT--AAACATTGTGAAATC--CTCGAT
MtPALM1   205 ATGTTCGTATATATATATATATAGGC-GA--AGATAGCTATGTTTTCAAAGTAGCTATAT


PsPALM1a  264 CTTGCGCTAGCC-ATACCC-T--------CTTCGACTTCGACTATGT---CTGGAAGGTT
PsPALM1b  242 GTTATCGAAGCT-TTATCAATGTGATAGATTGCAAGATCGAGAAAAAAATTTATGTTTTT
MtPALM1   262 GTTACTTCATCAGCTATCT-T--------TTGTTAATT-GTCT-TAT---ATGGAT-TCA


PsPALM1a  311 AGCAATTGCTTCGACTAGAATATCGGCAATTG--CTTCGACTTCGACTTCATCAGTTTC-
PsPALM1b  301 AATATGTGGTT-GGTTAGAGTATT-GTATTTA--CTTTG------CAGTGATGGATTGCA
MtPALM1   307 ATCAATTAATT--AATTGTCCAACGGGTTTTATAATTTGAATGCG-CGTAATGAGAGTTT


PsPALM1a  368 --TTCGTCAGCGTCATATGTCATCAATGGCT-TGTTAATTGGATCACCAGAATTCCAATC
PsPALM1b  351 AGTGCGAGAAAATTATTGGGGTTCAAAGTTTGTGT-TCTTATATAATCA-----------
MtPALM1   364 TATTTGTCTCTG-CAGTTG-CAGGGG-GGCTATGCATACT-GACAACAAGAAA-CTAACA


PsPALM1a  425 CCAGGCAGAATTTTCATCAATAACAATATCTCGACTCATCACGATCTTC-TCATTGATTG
PsPALM1b  399 ---TATAGTATTTGTTTTATTTATC-TAACCTTTT-TTATATAAAGCAAATAAATGT---
MtPALM1   419 GAATAGAGCATTAG--AGACCTACC-TAGCTTTGCCTTCTACCATTCTT-TCAACAT---


PsPALM1a  484 GATT-GAATAA-CATGTATGCTCCAGTTTTGTGATAT-TCTACCAGAATCAT--------
PsPALM1b  451 AAAAGTCATGAACATGTTGCATTAACATTTGATCT---TCCTCAAGATCGATGACGAGGA
MtPALM1   472 GATA-ATATAATTATTTTTTTTTTTTTTTTGTAAGATCTCAACAATAACAAT--------


PsPALM1a  533 -----AGGTTCAC---------ACTTGTCATCAAGCTTCC--TTTTACTCGCATCAGGAA
PsPALM1b  508 TCCGGATAATTAC---------TCTTCTCAGAGATCCAAC--ATTTATTCTT-CCAT-GT
MtPALM1   523 -----AGGTACAATGCAAGCAACTTAATCACTGATCATAAAAAAATATT-TT-AAAG-GC


PsPALM1a  577 CATGCTTGTA-ACAAATAGAGAAAAACACATTCAGTTGACTCACTGATGGACATTTGCCA
PsPALM1b  555 CGTG---ATG-ACG-ATGAAGAACAACAT-TTTACTTTACTATTTTGTGTAA--------
MtPALM1   575 CGTC---AAGTATT-TTGGAAACAAATAT-TTTATTAAAAGATTTTAC-TAA--------


PsPALM1a  636 CTCCATGCTTCTTCAGGGACCTTGTTTTTCAGCTTCTTGGTAGGGCACCTATTCA-GAAT
PsPALM1b  601 --ACA----------ATGTAGTTTGTAAACAA-------------------TTTA-ATTT
MtPALM1   621 --CCATTATCC-TTAGGGTAATTGTTAAGGAG-------------------TTCAAAAAT


PsPALM1a  695 ATAAATA-GCAAT-GGAAACAGATTCAC-------CCTATAAGAATTTTGGTAAATTCTT
PsPALM1b  629 ATTAATATTTCAT-GTAAACAATGTAATATATATATATATATATATA--TATATAT-AT-
MtPALM1   659 AGAAAATTTTCATTAAAAACA--ACAACCTTTTAACTTTCATAAAGT--TAAATAT---T


PsPALM1a  746 CTGCTT--CAGCATGCATCTCGGCATGTCCAAT-TTGGTCATGT--TTCTTCTTTCTTCT
PsPALM1b  684 ATATAT--ATATATATATATATATATATATATATATATATATATATATAT-CTTTCACCT
MtPALM1   712 CTACTTTTCAACA---ATTTCGTCTAATGCAGA-ATTTTTATATTTATCT-TTTTAACCA


PsPALM1a  801 -ATTCCATTATGTTGAATGGTGTAAGGAGCGGT-TACCTCAT-------GATCAAAACCA
PsPALM1b  741 CGGTTTGTTATAA-AATCGA-GAGGTAT--ATAGTGCTAGTTTGAAAAATATA-AAACTA
MtPALM1   767 -ACTCCCTTAGGG-CACTGA-TTAAC----ATT-T--TTCTT-------TATT-AAATAA


PsPALM1a  852 TATTCTGCACATAATTCTTCAAACATCTTTGAT-GTATATTCGCCACCTTCATCTGATC-
PsPALM1b  796 TT-GATGAGGTTGAACCCACGACCATTTAAAATAATCCCTTAGTTATTT--TT-AAATCG
MtPALM1   809 TT-GACGTAT--CAAAATTATAACATGGTTTAG-ATACATCAATTATTTA-AT-AAATCT


PsPALM1a  910 ---ACAAA-ACCTTGATCTTCTTTTCACTCTG---GTTCTCGACAA-GTATT---TTGAA
PsPALM1b  852 ACGGTAAATATCACACTTTTCATGTCATCCATTCATTACCTGGCATGGTAACAGGGTGAA
MtPALM1   863 A--AAAAA-AAAAAAATTTTTGTTTAAAACAG---TGACTTGA--A-GAAAT---AT-AT


PsPALM1a  959 TCTCTTAAAGATTGCAAATA-CTTCTTCCTTTCAC-TTGATCATATAGATCCAAAT-C-T
PsPALM1b  912 ATTCATTCACATTGGACATGACATGTGTTATTTATAGCGATGAGGGAC-TCCAATTACGT
MtPALM1   910 ATGTATAAATATTGCTTCTAATTTGTTTTT-TTCT-T---TTAAAAAG-GCCAAAA-GAA


PsPALM1a 1015 TTCGACTATACTCATCGACAAATGAGA-CAAAATACATGTTTCC-ACC-AATGGTATAAT
PsPALM1b  971 TCTGAG-----TCAAAGATTA-TCGGAGGACTTAGAATTCTAGAACAAAAAGGTTGATAG
MtPALM1   963 TATGA------TATAGGAAAA-TCGGAGTAGAAGACATACTTTAAATCAAATGATGGTAG


PsPALM1a 1072 CCTGAAATAGG-CCACATACATCTGAAT---ATACTACTTCTAG-TATGCAAGAGGATC-
PsPALM1b 1025 ATATCCATTGAACTAAGAAGTATCCCTTTAAGTAAATTATCTCTAGTGGTCATGT-ATCC
MtPALM1  1016 GTGTTTATAGG-TTATGGTCTTTGCAGT--GGTGATTTTTTTCG-GATGCCATAGAATTT


PsPALM1a 1126 ---TCATTGACATAG---TTAAAACGAAAA----ACTTAC--TGGATTGCTTTCTAACTA
PsPALM1b 1084 TACGAGTTGCTAGGAAAATGATAACTTTTGG---GGGGGC---AGCTATTCCAAATACTA
MtPALM1  1072 --CTTCTTCACATGTGTATTTTATTGTATCAGTGGCTGACATTGGATTTTCTTCCTATTG


PsPALM1a 1174 AGCAACCTTGATAG-AGTTTGTCGGGTATCTCATTAG-------TATACCAATTAACATA
PsPALM1b 1138 TGAAATACTAAATCCATTACCAGGACTTGATGATTACTAGGAAAATTGTGGGTGCGGAGA
MtPALM1  1130 TCTGATGTTGAGGTTACTTTGTACATCTTGTGATTATTGG-CTCTTTATCAATAAAAACA


PsPALM1a 1226 TCTTGAGT-AAT------CAGTG-TAAACAAATATATTCACAATTAATCGA-CGACTTGT
PsPALM1b 1198 AAGGAATA-CGTTGAGGCCACCGAGTAACAACCATCTTTAG-AGTGGCGCCACCAC-CAC
MtPALM1  1189 TGGAAAGAACATAAAAGACAACA-AAAACAAATAATTTAAA-TGAGATGGT-TCACATAT


PsPALM1a 1277 ATTCCA-CCG---AATCATACTTTAAT---TTAA---AAAAAA--ATATTTTAATGA-TT
PsPALM1b 1255 TTGTCTGACTAGGAAGAAGGAGTAAAGTCCCTAACGATTAAGAGAAGATTAGTTATTTGT
MtPALM1  1246 TTTTTAGTTT---AATAAGAGTTTAAGAATTTGA---TAGAAACGGAATTGCAAAGCCCA


PsPALM1a 1324 TAATAATTTATTTGTTTTTT-T-----GTTAGTTGAATAAAACT---ATTTTA-CACTAC
PsPALM1b 1315 GCATGCTCACACTAAAAAATACAAACTATTATC-CAGTTAAATCACAAACCCAAGACCGC
MtPALM1  1300 TAACTCTCTTTTCGTAGCATTTAGAACTTTATATTTGTAAAATG---ATTTCA-AACAA-


PsPALM1a 1374 TGCTAGTGCATA--TTCATTATATATA----TAT-ATATATATATATATATATATAATAT
PsPALM1b 1374 CACTCCCACCTACCCTCATCCCCCACAAAAACACTCTTTCTGCTAAGAGATAGAGAAAAG
MtPALM1  1355 --CTAGTATATA--TTGAATTAACGTGTG-ATGG-TTTTACATTTATTAACTCAGAAATT


PsPALM1a 1427 ATATATATATATATATA-TATATATATAATATATATATATATATTATATATATATATATA
PsPALM1b 1434 CTACAAAAATCTAACTTTGAGAGGAATCAAATCCACCCAGGG---ATCACTCCATAAATT
MtPALM1  1409 CTAAT-GAAAATAGGTAATGGAGTATTTATATTTGGG--------AAATTTTCATATTAA


PsPALM1a 1486 -TATATATATATATATATTAATTTAGTGATA----ATCTTTTAGAAGAAAACTGTTTATG
PsPALM1b 1491 -AGCGAGGAGACCAAAACCAACTTTCTTAACAACACTATCCTGAAATCAATTGATGGGTC
MtPALM1  1460 AAAAATAGGGAG-TGTAATAATTTTTTTTTT----TTTTTTTGAGGGGAAGGAGT--GTA


PsPALM1a 1541 CTCACTGTTGTTTCTCCTTACTG-TTTTT-TTTTCCGGCCTGTTTTGATGCTACCTG-CT
PsPALM1b 1550 CTCTCCTTGGCTACCTAGAGGAGATACCT-TTCTCCAC-GAAGGAGACGCGTAGCAAGAG
MtPALM1  1513 ATAAATTTTA--ATCAAAT--AG-TACCTTTTATCCAT-TATTTTTGTGAATATCAA-AT


PsPALM1a 1598 T-GTGGAGTGTTTGTAACTCTTGAGCTTTTATCTGAGAATGTTGGGCTTGGATTCAACTT
PsPALM1b 1608 AGATAACTCTGCCACCC--GAGACAGAAGTA-GTATGAATAACTTCATAC---CTAGCAT
MtPALM1  1566 TTATTTTTTTACTAAAAATGATAAGGATCCATCTTTGAACTATTTTTTAGAGGTCCACAC


PsPALM1a 1657 TTTAAATATATTTTGTTGATTAAAAAGATTATT-----AATTAATTATACAATAACTTGA
PsPALM1b 1662 ATAGGATATCCCAC--CATAAACTTGCATCACCCG--AAATCAATTG-----CCACCCCA
MtPALM1  1626 ATTAGATGCACCCT-TAAAAAAATTATATTAGCATATGAATAAAGTA----ATAAACTAA


PsPALM1a 1712 AATTTTAATGACT-TAGTACGTATTTATGGAATGTTAAC--TTATGACAAGTTAATATGT
PsPALM1b 1713 TTTACTTAACAAA----GTCAGAC-TAA-CCATCCAAAGATCACGAACACCTAAGCCCCC
MtPALM1  1681 TTTATGAATTAATGTTATTCGTAT-TATGTTATTTTAAGAGGTAGAACAAGTTGGGTCAA


PsPALM1a 1769 AT-ATATTT-TTTTG---GTTACATATATAACAAGTTAATTTA-TCAA-GTTTTAAAATA
PsPALM1b 1767 ACACCTCTTGGTTTTGCGAACA-TCAGATCAC---CTACCCCAAACA--ACCTTAGAG-C
MtPALM1  1740 ACAATATTT-TTTTT--AAAGATTTTGATTACTATATATCTTA-GAAGCATTTTAATG-A


PsPALM1a 1822 TGGATTTA--AT-------------------TGGGATATA---AACATTAATAAAAGGAA
PsPALM1b 1820 CTCATCTAACACCACCCCACAAAAATCTTCTTTGAATTCTCACAATCTTCTTTCAAACCG
MtPALM1  1795 AATATTTATTAT-------------------TTTAAAATT--CATTTTTATTAAAAATAG


PsPALM1a 1858 GAAAAATAAATAATCTACACTGATATTT---ATAGAGTAATAATTTTTTTAT--TATT-C
PsPALM1b 1880 TA-----------ATAGG---CATCTTTATAAAAGACAAAAAGAAGATAGAAATAGCATT
MtPALM1  1834 TAAATGTTTAT-ATTTGGAAGAATTTTG---AATGATTCATATTAGATTGACCCTATA-T


PsPALM1a 1912 AAACATGCACTTATAG-AA--CT------CATATAATTGAATAAAATAATTA-------A
PsPALM1b 1926 AAGAACATAATTG-AG-AAACCGCCTAGTGATAAGATTAATTAAA--AATTC-------C
MtPALM1  1889 AGGCAGTTACTTTTATGTA--TT------TCTACT-TTTTTTAGGA-GATGTATATTTAT


PsPALM1a 1956 TATT-AAAATAAA--ATATAATTAA---TATTATTATTCA-AT---TCAATTACT-AATC
PsPALM1b 1975 TATGTAAAGGACAATCGAGAATTCG--CCCCTACTAAAAAAAC---CTAAATACTTAAAT
MtPALM1  1939 TTTGTATAG-CCTTTTTACATTTTGAATGCTTAATGTCAAAATTAAGGACATATTAAACT


PsPALM1a 2005 AATA--AAAACTTTTA-ATAAATAACTAA-TATTATT-GATTATAAAGGTATTTACCAAC
PsPALM1b 2030 GATA--GAGACTCTCT-TT-AACAGTGAA-TAAAATCACATTTTAAAAATATTTGCCAAC
MtPALM1  1998 TATATTGCGATTTGCAACT-AATTAAAAATTAAAAAG-AGGGTGAGGAATATT-GAAGAT


PsPALM1a 2060 ATGATGTTGA-AATCTA--ATTAAATAATAAAA-TATTTCAACTTGTTTAACCTTTTTAC
PsPALM1b 2085 ATAATCTCAAAAATCTC--AAAAATCTCAAAAA-TCTTT---------------------
MtPALM1  2055 TTAAA-ACAATGGTGTGCTTTCCCCCTCAAAAAATAATTG--------------------


PsPALM1a 2116 ATTTTGAATATCTTATGTCAAATTTAATAAACAGTTTGCCTTCAATGGCGTAGATGCCAA
PsPALM1b 2121 ---------------TAA---TCCTGAT---A-GATACA---CAAAGATA-AAATCTTTT
MtPALM1  2094 ---------------TGT---GCTTGAT---GAGTTATATTTCAATGACGTAGATGTCAA


PsPALM1a 2176 AGGAAATAAT-TGTCTTTATTAAGC--TAGCTAGTTGTATGAGATGACAATTTATTAAAA
PsPALM1b 2155 AATAAACAACTAA--TATAATTTGCAAAACATAAT-GT-TGAAAT-CTAA----TTAAAT
MtPALM1  2133 AGGAGATGATTTGTCTATATTAAGCAATAGCTAGTTGTATGTGCT-TCAA----TTATAA


PsPALM1a 2233 CCA-------------------------------AATATAATTGTC-TTTTACCATTATT
PsPALM1b 2206 GATAAAATATTTCAACATGTTTACCCTTTTTACATATTTGAATATC-TTATGTAAATACT
MtPALM1  2188 CCT-------------------------------ATAGTATTTTTTTATTTACCATTATT


PsPALM1a 2261 AA---------------------------AATACTACTTTTCTTTTTTAATATAA---CC
PsPALM1b 2265 AA---------------------------TAAATAGTTTGCCTTCAATGGCGTAGATGCC
MtPALM1  2217 AGTATAATATATCCTTTTTTTTTTTTGAGGGAGTAGTATAGTATAATATATCTAG-TGTC


PsPALM1a 2291 -AGTAACATGTAGTGTCA--ATCTCACTATG-TTCCAACTTCAAATTTATTAATGGGTGA
PsPALM1b 2298 -AAAGGAAAGAATTGTCTTTATTAAGCTATG-ATCCAACTTCAAATTTATTAATGGGTGA
MtPALM1  2276 AATATGTTCCAATCCAG—CAGCTATCTATCTAGCTTACTTAAAATTTATTAATTGGTGA
```

PsPALM1a 2347 TACATTCGATTTGATGTTTTAAAGTTTAACTT**TGTCTT**C---CTAAA-----TTAATAAT
PsPALM1b 2356 TACATTCGATTTGTTGTT-TAAAGTTTAACTC**TGTCTT**C---CTAAA-----TTAATAAT
MtPALM1 2334 TACATTCGATTTGATGTTTTAAAGCATGACTT**TGTCTT**GACCTTAAATATTATTAATTAT


PsPALM1a 2399 ATTCTCTA-CTATATTTAAATAAGCAATTATTTTAATCTTCATTTTCATCCCCC-----A
PsPALM1b 2407 ATCCTCTA-GTATATTTAAATAAGCAATTATTTTAATCTTCATTTTCATCCCCC-----A
MtPALM1 2394 ATTCACCCCCCCTATTT-AATAACGAATTATATTTATCTCCAATTTCATCCCCCACCCCA


PsPALM1a 2453 TCACCAATATCTTTCTATCTTTCTATCTTAAA**TGTCAGCGACA**GTACCATGCCTACAGAT
PsPALM1b 2461 TCACCAATATCT--------TTCTATCTTAAA**TGTCAGTGACA**GTACCATGGCTACAGAT
MtPALM1 2453 TTACCAATATCTATCTATCTTTCTATCTTAAA**TGTCAGTGACA**GTACCATGGCTACAGAT

```
PsPALM1a 2513 ATTT---------CCATCACTACTACTCAGATCCAAAATTCATCACAATCACAATCTCAA
PsPALM1b 2513 ATTGCCCTTCTTTCCATGACTACTACTCAGATCCAAAATTCATCACAATCACAAT-----
MtPALM1  2513 ATTGGCCTTCTTTCCA---ATATGACTCAGATCCAAAAATCATCACAATCACAATCTCAA


PsPALM1a 2564 T---CTCAACCAAACCCTAACATCATCACCACC------ACCACACCATCATCATCAACT
PsPALM1b 2568 ----------CAAACCCTAACACCACCAGCACC------ACCACACCATCACCATCAACT
MtPALM1  2570 CAACACCAACCAAACCCTAATTCCAACACCACTACACCCCCCTCACCATCTTCATCAACT


PsPALM1a 2615 TGGATGTGGAACCCTAAACAACAACAACATCAAGAACAAGAAGATGAAGATTCATGGGAG
PsPALM1b 2612 TGGATGTGGAACCCTAAACAACAACAACATCAAGAACAAGAAGATGAAGATTCATGGGAG
MtPALM1  2630 TGGATGTGGAACCCTAAACAAC------ACCAAGAACAAGAAGATGAAGATTCATGGGAG


PsPALM1a 2675 GTAAGAGCTTTTGCAGAAGACACAAGGAACATGATGAACACAACGTGGCCACCAAGATTC
PsPALM1b 2672 GTAAGAGCTTTTGCAGAAGACACAAGGAACATGATGAACACAACGTGGCCACCAAGATCC
MtPALM1  2684 GTAAGGGCTTTTGCTGAAGACACAAGGAATATTATGAACACAACATGGCCACCAAGGTCC


PsPALM1a 2735 TACACCTGCACTTTTTGTAGAAGAGAGTTCCGGTCAGCTCAAGCTCTTGGCGGTCATATG
PsPALM1b 2732 TACACCTGCACTTTTTGTAGAAGAGAGTTTCGGTCAGCTCAAGCTCTTGGCGGTCATATG
MtPALM1  2744 TACACCTGTACCTTTTGCAGAAGAGAGTTCCGGTCAGCTCAAGCTCTTGGTGGTCACATG


PsPALM1a 2795 AATGTCCACCGCCGCGACCGTGCTCGTCTCCATCAAAATCAACCACCGTTAAACTCCTCC
PsPALM1b 2792 AACGTCCACCGCCGTGACCGTGCTCGTCTCCATCAAAATCAACCACCGTTAAATTCCTCC
MtPALM1  2804 AATGTTCACCGCCGCGACCGTGCTCGTCTCCATCAAACCCAACCACCGTTAAATTCTTC-


PsPALM1a 2855 TCTCATCATCCTTCTTCTCCATTCATACATATCCCTCCTCAAGAGCTTGT---TGATGCT
PsPALM1b 2852 TCTCATCATCCTTCTTCTCCGTTCATACATATCCCTCCTCAAGAGCTTGT---TAATGCT
MtPALM1  2863 -----TCATCCTTCATCACCATTCATAAATATTCCTCCACAAGACCTTGTTGCTAATGCT


PsPALM1a 2912 GGATTGTGCCTTTTTTACCATTCACCAAACCCTAATAT---------TTCTTCCTTCAAT
PsPALM1b 2909 GGATTGTGCCTTTTTTACCATTCACCAAACCCTAATAT---------TTCTTCCTTCAAT
MtPALM1  2918 GGATTGTGCCTTCTTTACCATTTACCAAACCCTAATAATAATGCTTTTGCTTCCTTCAAT


PsPALM1a 2963 GATTC------TAATGGAGAATCTCCTTCAACTTTTCTCTCTATATCATCAT---CATCT
PsPALM1b 2960 GATTC------TAATGGAGAATCTCCTTCAACTTTTCTCTCTATCTCATCAT---CATCT
MtPALM1  2978 AGTTCTAGTCCTAATGGAGAATCTCCTTCTACTTTTCTCTCAATCTCATCATCAACATCT


PsPALM1a 3014 TATCCAACAAACAACTTGATGATGCAAATGCAAATGCAAACTTGTTCTCCTCCATCTTTT
PsPALM1b 3011 TATCCAACAAACAACTTCATGATGCACA------TGCAACCTTGTTCTCCTCCATCTTTT
MtPALM1  3038 TATCCTCCAAATAATTTGATGATGCAAA------TGCAAGCTTGTTC---TCCATCTTTT


PsPALM1a 3074 CATTTTCAAG---CTAATTCAGCTAGAAATTTGATTAACAATAGCATATCTTCTTTTTCT
PsPALM1b 3065 CATTTTCAGG---CTAATTCAGCTAGAAATTTGATTAACAATAGCATCTCTTCTTTTTCT
MtPALM1  3089 AATTTTCAAGTGGATAATTCAGCTAGG---TTGATCAACAATAGCATTTCTTCTTTTTCT


PsPALM1a 3131 AACAAAC------------CTGCTATCTGCACCTCCATTAATGATAAGGTTCATGAAATT
PsPALM1b 3122 AACAAAC------------CTACTATCTGCACCTCCATTGATAATAAGGTTCATGAAATT
MtPALM1  3146 AGCAAAGTGGACCAGCAACATGCTACTTGCACCTCCATTGATGATAATGGTCATGAAATT
```

```
PsPALM1a 3179 GAAGAACTCGATCTTGAGCTACGTTTGGGGAACAAGCCATCACCGGCATGAAAAATCTAT
PsPALM1b 3170 GAAGAACTCGATCTTGAGCTACGTTTGGGGAACAAGCCATCACCGGCATGAAAAATCTAT
MtPALM1  3206 GAAGAGCTTGATCTTGAGCTCCGTTTAGGGAACAAGCCAACACCAACTTGAAAAGG-CAT


PsPALM1a 3239 AGCTACATCATAGATATATAGATGGATGAATATTTATTTATATTTGAAGTTATTAATTCA
PsPALM1b 3230 AGCTACATCATAGATATATAGGTGGATGAATTTTTATTTATATTTGAAGTTATTAATTCA
MtPALM1  3265 AG-------ATAGATATTTATAT--TTGAAGTTTGAACCATATATG--GTGGTTAATTGA


PsPALM1a 3299 TTTTAGTGTGTATGAAGAATATT-GAATTATAAATTCCTAAAAGTATTCTTATAT-ATTA
PsPALM1b 3290 TTTTAGCGTGTATGAAAAATATT-CAATTATAAATTCCTACAAGTACTCTTATAT-ATTA
MtPALM1  3314 TTTTAGTGTATGTGAAGAAAATTTAAATTCTAAATTCCAACAAGTACTTTTAAATGAATT


PsPALM1a 3357 ATTTGTTC--TCTGTCTTTTGCAGCGTGTAGTGAAGAAGAAATTAGTGTCATGTAATTTT
PsPALM1b 3348 ACTTGT----TCTCACTTTTGCAGTGTGTAGTGAAGAAGAAATTAGTGTCCTGTAATTTT
MtPALM1  3374 ATTTGCTCGATCTCTCTTTTTCAGTGTGTGG---AGAAGAA---AGTGTACTGTAATT-T


PsPALM1a 3415 CTTAATGATGGAGACACCATGTCCAGCTA------TACATGAAATTGATAGCACTATAAA
PsPALM1b 3404 CTTAATGATTGAGACACCATGTCCAGCTAGCTTTTTTTATGAAATTGATAGGACTATAAA
MtPALM1  3427 GTTGATGATTGAGACACCATGTCCAGCTTT-TACATAGCTGAAATTGATAGGACTATG--


PsPALM1a 3469 TGGATTTTGTTTTTGGTTGGGGTTTTTGGGGTTAATTTCAATAA-----------TTTAT
PsPALM1b 3464 TGGATTTTGTTTTTGGTTGGGGTTTTGGGGGTTAATTTCAATAA-----------TTCAA
MtPALM1  3484 -GGATTTATTTTTTGGTTGGGGCTCTAGGGATTAATTTCAATAAAATTTCAATAATTTAT


PsPALM1a 3518 TGATTTTTTTGGTTGATGGAATTTATTGTTATA-TGA-----------------------
PsPALM1b 3513 CATTTTTTGTGGTTGATTGAATTTATTATTATG-TGACTG-------GTTTTGTGGCTAT
MtPALM1  3543 TGTTTCTTTGGGTTGATTGATTTTATTAATATGATGAGTGTTTAATTGTTTTGTAACTAT


PsPALM1a 3554 --TTATGAATTGCAAGTATATTAATTGTAATGATGGAAAAAGATTAGGGTTTTGAAGAAG
PsPALM1b 3565 GATTCTGAATTGCAAGTA--GTAATTGTAATGATAAAAAAAAATTAGGGTTTTGAAGAAG
MtPALM1  3603 GAAGGTGAACAAGAAGGA--GCAATT--AGTTA-------------TTCATTTGGGAAGG


PsPALM1a 3612 GCACCCAAAAGGTTGCAG---------TGAAGTGTCATTTTCATGTTTACACACAAATAT
PsPALM1b 3623 ACATTGAAAAGGTTGCAG---------TGAAGTGTCATTTTCTTGTTTACACACAAATAT
MtPALM1  3646 GAATGGGAAAGGAGGAGGAGGAGAGAAAGAAGAGGATTTTTCAAAGAAAAATTAAAATGT


PsPALM1a 3663 ATGTTTCTGGGCATACATTTCCGTGGGGATTATTATTATTTTCATCT-TTTTTTCCTAAA
PsPALM1b 3674 ATGTTTCTGGGCATACATTTCCGTGGGGATTATTATTATTT-------------------
MtPALM1  3706 AAGTTGGTA------------------TATCACTATTTTTTTAAAGATATTTCAAGTAGA


PsPALM1a 3722 CACTAAT-A
PsPALM1b 3715 ---TCATCT
MtPALM1  3748 AACTAATGA


(B)

PsPALM1a    1 ATAAAAACTTTTAATAAATAACTAATATTATTGATTATAAAGGTATTTACCAACATGATG
PsPALM1b    1 ATAAAATCTTTTAATAAACAACTAATATAATTTGCAA--------------AACATAATG


PsPALM1a   61 TTGAAATCTAATTAAATAATAAAATATTTCAACTTGTTTAACCTTTTTACAT-TTTGAAT
PsPALM1b   47 TTGAAATCTAATTAAATGATAAAATATTTCAACATGTTTACCCTTTTTACATATTTGAAT


PsPALM1a  120 ATCTTATGTCAAATTTAATAAACAGTTTGCCTTCAATGGCGTAGATGCCAAAGGAAATAA
PsPALM1b  107 ATCTTATGTAAATACTAATAAATAGTTTGCCTTCAATGGCGTAGATGCCAAAGGAAAGAA


PsPALM1a  180 TTGTCTTTATTAAGCTAGCTAGTTGTATGAGATGACAATTTATTAAAACCAAATATAATT
PsPALM1b  167 TTGTCTTTATTAAGCTA-------------------------------------------


PsPALM1a  240 GTCTTTTACCATTATTAAAATACTACTTTTCTTTTTTAATATAACCAGTAACATGTAGTG
PsPALM1b  184 ------------------------------------------------------------


PsPALM1a  300 TCAATCTCACTATGTTCCAACTTCAAATTTATTAATGGGTGATACATTCGATTTGATGTT
PsPALM1b  184 ------------TGATCCAACTTCAAATTTATTAATGGGTGATACATTCGATTTGTTGTT


PsPALM1a  360 TTAAAGTTTAACTTTGTCTTCCTAAATTAATAATATTCTCTACTATATTTAAATAAGCAA
PsPALM1b  232 T-AAAGTTTAACTCTGTCTTCCTAAATTAATAATATCCTCTAGTATATTTAAATAAGCAA


PsPALM1a  420 TTATTTTAATCTTCATTTTCATCCCCCATCACCAATATCTTTCTATCTTTCTATCTTAAA
PsPALM1b  291 TTATTTTAATCTTCATTTTCATCCCCCATCACCAATATCTTTCTATCTT--------AAA


PsPALM1a  480 TGTCAGCGACAGTACCATG
PsPALM1b  343 TGTCAGTGACAGTACCATG
```
